# Hypoosmolarity inhibits archaeal ammonia oxidation

**DOI:** 10.1101/2025.07.07.663500

**Authors:** Joo-Han Gwak, Adebisi Olabisi, Ui-Ju Lee, Christiana Abiola, Seongjun Lee, Hackwon Do, Yun Ji Choi, Jay-Jung Lee, Man-Young Jung, Nico Jehmlich, Martin von Bergen, Michael Wagner, Samuel Imisi Awala, Zhe-Xue Quan, Sung-Keun Rhee

## Abstract

Salinity strongly influences the physiology and distribution of nitrifying microorganisms, yet the effects of low salinity on them remain understudied. This study investigates the impact of hypoosmolarity on different groups of ammonia oxidizers in soil and lake environments, as well as in pure culture isolates. In soil microcosms amended with ammonium, at low salinity levels (∼120 μS/cm), comparable to values commonly found in pristine terrestrial and aquatic environments, the abundance of ammonia-oxidizing bacteria (AOB), dominated by *Nitrosomonas oligotropha*, significantly increased. In contrast, the growth of ammonia-oxidizing archaea (AOA), dominated by “*Ca.* Nitrosotenuis” of the *Nitrosopumilaceae* family, was stimulated by high salinity (∼760 μS/cm). In ammonium-fed lake microcosms, the abundance of AOB, dominated by *N. oligotropha,* significantly increased under both low (∼170 μS/cm) and high salinity (∼850 μS/cm) conditions. In the presence of allylthiourea, a bacterial nitrification inhibitor, AOA were sensitive to low salinity in both soil and lake microcosms. Consistently, pure culture studies revealed marked growth inhibition of AOA, especially members of *Nitrosopumilaceae*, under hypoosmolarity, unlike AOB and complete ammonia oxidizer (comammox) strains. Comparative genomic analyses with AOB and comammox, along with transcriptomic studies, suggested that the sensitivity of AOA to hypoosmolarity stress was possibly due to a lack of sophisticated osmoregulatory transport systems and their S-layer cell wall structure. Overall, this study highlights hypoosmolarity as a key factor shaping the ecological niches and distribution of ammonia oxidizers, as well as nitrification activities, in terrestrial and aquatic environments that are increasingly affected by intensified water cycles due to climate change.

## Introduction

Ammonia oxidation is the first and often rate-limiting step of nitrification in the nitrogen cycle [1]. This process is mostly driven by chemolithoautotrophic ammonia oxidizers, which conserve energy by oxidizing ammonia (NH3) to either nitrite (NO_2_ ^-^) or nitrate (NO_3_ ^-^) [1–4]. As climate change accelerates the water cycle [5, 6], fluctuations in salinity across terrestrial and aquatic environments are becoming more pronounced due to shifts in precipitation patterns and more frequent extreme weather events such as flooding and drought [7, 8]. These changes, in turn, alter microbial habitats by disrupting osmotic balance and ion homeostasis [9, 10], both of which are critical for the stability of microbial communities that mediate the nitrification process [11–13]. Despite the widespread occurrence of hypoosmotic conditions in terrestrial and freshwater environments [14, 15], little is known about how microorganisms, particularly ammonia oxidizers, adapt to chronic hypoosmotic stress. Investigating their responses under environmentally relevant salinity conditions is essential for predicting shifts in nitrogen cycling in terrestrial and aquatic ecosystems influenced by global water dynamics.

Climate change, driven by global warming, is significantly intensifying the Earth’s water cycle [5, 6]. This intensification leads to localized desertification and flooding events that occur more frequently across the globe, highlighting the growing imbalance in water distribution [7, 8]. Projections indicate that by 2100, the planet will experience a marked increase in heavy precipitation events [16, 17], which will substantially alter hydrologic dynamics in soil systems. For instance, in the United Kingdom, soil water saturation levels are expected to rise during winter, with prolonged periods of saturation [18]. These precipitation changes are likely to have far-reaching effects on various environmental parameters, particularly salinity. As an important hydrologic factor, salinity is highly sensitive to fluctuations in precipitation patterns. Generally, increased precipitation tends to dilute and thus decrease the salinity of soil pore waters, surface waters (such as lakes and rivers), and groundwater in aquifers. In addition to climate-driven changes, anthropogenic factors such as irrigation and fertilization also significantly influence soil salinity, often resulting in elevated salt concentrations [19–21]. These agricultural practices can alter the ionic composition of soil solutions and may exacerbate or counterbalance the effects of natural precipitation patterns. Moreover, these salinity changes have cascading effects on other environmental properties, including pH, redox potential, and metal availability [7, 22–24].

Shifts in salinity are key abiotic factors influencing microbial physiology. The normal functioning of microbial cells relies heavily on the regulation of intracellular ion concentrations [9]. Osmoregulation, the process by which cells maintain water and salt balance, requires considerable energy and is fundamental for cellular survival under both hyperosmotic and hypoosmotic stress conditions. Consequently, variations in salinity can significantly influence the physiology and ecology of microorganisms [25, 26], as well as other organisms inhabiting terrestrial and aquatic ecosystems. These shifts in microbial physiology, driven by changes in salinity, may subsequently impact key biogeochemical processes, potentially disrupting nutrient cycling, modifying microbial production and consumption of greenhouse gases, and altering overall ecosystem function.

Nitrification, the sequential aerobic oxidation of reduced nitrogen compounds primarily NH3 to NO ^-^ via NO_2_ ^-^, is considered pivotal in the global biogeochemical nitrogen cycle on Earth. Furthermore, ammonia oxidation is a key process that contributes to the release of nitrous oxide (N2O), a potent greenhouse gas and ozone-depleting substance [27, 28]. This process is microbially executed by different yet co-occurring groups of chemolithoautotrophic ammonia oxidizers, specifically ammonia-oxidizing archaea (AOA) and ammonia-oxidizing bacteria (AOB), which can oxidize NH3 to NO ^-^ using it as their sole energy-conserving substrate [1, 2, 29]. A new group of bacterial ammonia oxidizers, which perform complete ammonia oxidation (comammox; CMX) of NH3 to NO ^-^, was recently discovered [3, 4]. This group consists of clades A and B, which exhibit different distributions depending on habitat [4, 30, 31]. AOA, AOB and CMX differ in their kinetic properties and N2O-yields [32–36]. Regardless of the type of ammonia oxidizer, nitrification results in fertilizer loss and is therefore commonly targeted with nitrification inhibitors in agricultural systems. However, the sensitivity of AOA, AOB, and CMX to such inhibitors can vary [37], suggesting that salinity-induced shifts in ammonia-oxidizing community composition could influence both the rate of N2O emissions and the effectiveness of nitrogen retention strategies in managed soils.

Previous studies on the impact of salinity on nitrification have mostly focused on hyperosmolarity, as high salinity is often a serious concern in agricultural and environmental contexts [11, 13, 38–42]. In contrast, salinity below 350 μS/cm has been associated with reduced microbial diversity in a freshwater microcosm study [43], suggesting this level as an ecologically meaningful threshold for defining hypoosmotic stress [44, 45]. In pristine environments, salinity— typically assessed via electrical conductivity (EC)—is often well below this threshold. In lakes, the 25th percentile EC is approximately 100 μS/cm, the median is 227 μS/cm, and the 75th percentile is 427 μS/cm (Supplementary Fig. S1) [14], highlighting the prevalence of hypoosmotic conditions. Similarly, in soils, EC values derived from soil water extracts show a median of 130 μS/cm and a 75th percentile of 410 μS/cm (Supplementary Fig. S1) [15]. These salinity levels are substantially lower than those of typical laboratory mineral media [33, 46–48], such as the artificial freshwater medium (AFM; ∼3,600 μS/cm) [35], which is commonly used to cultivate and study various microorganisms, including terrestrial and freshwater ammonia oxidizers.

Investigating the impact of environmentally relevant hypoosmotic conditions on ammonia oxidizer activity is vital for understanding current nitrogen cycling processes and predicting future changes in nitrogen cycling dynamics due to shifts in global water cycles [49, 50]. While most laboratory studies on hypoosmolarity have focused on the microbial cell responses to rapid and acute decreases in salinity (i.e., hypoosmotic shock) under laboratory conditions [51–54], few have examined the adaptation of microorganisms to ‘chronic’ hypoosmotic stress [55–58]. In this study, we investigated the effect of chronic hypoosmolarity on ammonia oxidizer activity in soils and lake waters. The activity of AOA, particularly those within the *Nitrosopumilaceae* family (hereafter referred to as AOA-NpF) [59], was sensitive to hypoosmolarity, a finding further supported by pure culture experiments. Overall, we propose that hypoosmolarity serves as a potential key environmental factor contributing to the niche differentiation and abundance of ammonia oxidizers in low-salinity terrestrial and aquatic environments.

## Materials and Methods

### Environmental salinity and media composition

Electrical conductivity values (EC; μS/cm), as a measure of salinity, were obtained from global datasets of lakes [14] and soils [15] to characterize typical environmental salinity ranges. To ensure relevance, the datasets were filtered to include only non-coastal lakes and soils with non-zero EC values, as zero entries may reflect conditional imputation in the original dataset [15, 60, 61]. Subsequently, EC percentiles were calculated to identify representative salinity levels typically observed in these environments. In combination with data on natural freshwater salt composition [62], these EC values informed the design of a mineral water medium (MWM) intended to mimic the ionic compositions of natural freshwaters (Supplementary Table S1). Varying salinity levels were achieved by preparing different strengths of MWM through the adjustment of monovalent and divalent cation concentrations.

For microcosm experiments, NH4Cl was used as a substrate. KH2PO4, NaHCO3, HEPES, pyruvic acid, trace elements, and ferric sodium EDTA were added as supplements depending on experimental conditions, with concentrations detailed in Table S1. After all supplements were added, the EC of 1× MWM was measured at 128 μS/cm, corresponding to the lower EC range observed in lake waters and soil environments. In contrast, 10× MWM had an EC of 795 μS/cm, representing the higher EC range observed in lentic waters and soils, and was used for experiments requiring elevated salinity conditions (Supplementary Fig. S1). MWM was used to examine the effects of salinity on ammonia oxidizers in both microcosm and pure culture experiments. Artificial freshwater medium (AFM) [63], previously described and with an EC of 3,540 μS/cm, served as a control.

### Microcosm experiments

Soil samples (top 30 cm) were collected from grassland and agricultural fields. The agricultural field was fertilized annually with oil cake (N:P:K = 4:2.1:1; 70% organic matter content) and potassium chloride, providing approximately 190 kg N ha^-1^ year^-1^, 48.8 kg P ha^-1^ year^-1^, and 123.7 kg K ha^-1^ year^-1^, split into three applications from May to September. Metadata, including sampling location, date, depth, EC, pH, and nutrient levels, are provided in Supplementary Table S2. The samples were sieved through a 4-mm mesh and stored at 4°C until use. Soil slurry microcosms were set up in 250-ml polystyrene culture flasks, each containing 3 g of soil mixed with 150 ml of 0×, 1×, or 10× MWM, with triplicate samples for each condition. The intrinsic salinity contribution from soil samples was negligible. Lake water samples were collected at 20 m depth, corresponding to the beginning of the aphotic zone, from Lake Soyang and Lake Daecheong, with sampling details provided in Supplementary Table S2. Lake water microcosms were prepared using unfiltered fresh lake water alone or supplemented with 10× MWM basic salts, in triplicate for each condition.

NH4Cl (25 μM or 100 μM) was added as a substrate to the soil and lake microcosms, respectively. All microcosms were incubated at 25°C in the dark with intermittent inversion. Ammonia, nitrite, and nitrate concentrations were measured to monitor nitrification activity, following previously described methods [63]. To specifically monitor nitrification by AOA, a control microcosm was treated with allylthiourea (ATU; 50 μM) to selectively inhibit AOB and CMX [64–66]. Once ammonium oxidation was complete, all microcosms were harvested by filtration using 0.1-μm pore-size mixed cellulose ester filters (Advantec, Japan) and stored at -80°C before DNA extraction.

### DNA Extraction, qPCR, and 16S rRNA gene amplicon sequencing

Genomic DNA was extracted from filters that had been ground in liquid nitrogen using the Soil DNA Kit (GeneAll, Korea) according to the manufacturer’s instructions. DNA concentration and integrity were assessed using a Qubit™ 4 Fluorometer (Thermo Fisher Scientific, USA) and by agarose gel electrophoresis. Extracted DNA samples were stored at -80°C for downstream applications. Quantitative PCR (qPCR) was conducted using specific primers targeting AOA, AOB, and CMX groups, with sequences and conditions provided in Supplementary Table S3.

The hypervariable V4–V5 region of the 16S rRNA gene was amplified using the primer pair 515F/926R, along with sample-specific indexing adapters (Nextera XT Index Kit), following the manufacturer’s instructions. The amplification consisted of an initial denaturation at 95°C for 3 minutes, followed by 25 cycles of 95°C for 45 seconds, 50°C for 45 seconds, and 72°C for 90 seconds, and a final extension at 72°C for 5 minutes. Amplicons were purified using the Labopass purification kit (Cosmo Genetech, South Korea), and their concentration and quality were assessed as described above. Sequencing was performed on the MiSeq platform (Illumina) with 300-bp paired-end reads by LabGenomics Inc., Korea. Raw sequence reads were quality-checked using FastQC (v0.11.8) [67], and adapter trimming and quality filtering were performed using Cutadapt (v4.9) [68], as integrated into the QIIME2 pipeline (qiime2-amplicon-2024.10) [69]. Reads were demultiplexed in QIIME2 and processed to generate amplicon sequence variants (ASVs) using the DADA2 package (v1.20.0) [70] within R. ASVs associated with chloroplasts, mitochondria, and singletons were filtered out before downstream analysis. Taxonomic classification of ASVs was performed using the Greengenes2 database (2022.10) [71].

### Phylogenetic analysis

ASVs of 16S rRNA genes were aligned using MAFFT (v7.526) [72] with L-INS-i algorithm based on structurally curated seed alignments of bacterial and archaeal 16S rRNA gene sequences retrieved from the Comparative RNA Website (as of July 2024) [73]. Phylogenetic relationships were inferred using the maximum-likelihood method implemented in IQ-TREE (v2.3.6) [74], with ModelFinder (-m MFP) [75]. The phylogenetic tree was visualized using iTOL (v6.9.1) [76].

### Effect of hypoosmotic stress on ammonia oxidizer strains

The effect of hypoosmotic stress on growth was tested across a range of salinity levels (0–128× MWM) with four AOA strains—two from AOA-NpF: *Nitrosarchaeum koreense* MY1, isolated from agricultural soil in Korea [77], and “*Candidatus* Nitrosotenuis chungbukensis” MY2, isolated from a deep oligotrophic soil horizon [78]; and two from the *Nitrososphaeraceae* family (hereafter referred to as AOA-NsF): *Nitrososphaera viennensis* EN76, isolated from garden soil [79], and “*Candidatus* Nitrosocosmicus oleophilus” MY3, isolated from coal tar-contaminated sediment [80]; as well as one ammonia-oxidizing bacterial strain, *Nitrosomonas europaea* ATCC 19718, isolated from soil [81], and the only complete ammonia-oxidizing bacterial strain available in pure culture, “*Candidatus* Nitrospira inopinata” ENR4, isolated from a biofilm in a hot water pipe of an oil exploration well [4]. Cultures were incubated under oxic conditions with ambient air, without shaking, and in the dark. The incubation temperatures were set to 25°C for *N. koreense* MY1 and *N. europaea* ATCC 19718; 30 °C for “*Ca*. N. chungbukensis” MY2 and “*Ca*. N. oleophilus” MY3; 37 °C for “*Ca.* N. inopinata” ENR4; and 42 °C for *N. viennensis* EN76. The specific growth rate (μ) was calculated by determining the slope of the log-transformed nitrite concentration over time, using the equation μ = (ln*N*1 − ln*N*0)/(*t*1 − *t*0), where (ln*N*1 − ln*N*0) represents the change in the natural logarithm of nitrite concentration, and (*t*1 − *t*0) represents the change in time.

### Statistical analysis

To evaluate the abundance of ammonia oxidizers under different microcosm conditions, quantification based on qPCR data was analyzed using a two-way analysis of variance (ANOVA), with treatment conditions and types of ammonia oxidizers as independent factors. Tukey’s test was applied to identify pairwise differences between treatment levels. For 16S rRNA gene amplicon data, the relative abundances were square-root transformed and subjected to two-way ANOVA, followed by Tukey’s test, as described above.

### Comparative genomic analysis

To investigate the genomic repertoire of osmoregulatory systems in ammonia oxidizers, a total of 463 high-quality genomes (completeness >90%, contamination <10%, assessed by CheckM [82]) were retrieved from the NCBI database (Supplementary Dataset S1), including 266 AOA, 179 AOB, and 18 CMX genomes. The presence or absence of osmoregulatory genes was determined based on KEGG Orthology (KO) annotations using KofamScan [83] and BlastKOALA [84], both with default parameters. Phylogenomic relationships among genomes were inferred following the Anvi’o phylogenomics workflow [85], using IQ-TREE (v2.3.6) [74] with the LG+G+F model. The resulting phylogenomic tree was visualized in iTOL (v6.9.1) [76].

### Transcriptomic and proteomic analysis

To assess the differential expression of genes under varying salinity conditions, 1× and 10× MWM — representing low and high salinity in the environments — were used. Representative AOA strains from AOA-NpF (“*Ca.* N. chungbukensis” MY2) and AOA-NsF (*N. viennensis* EN76) were prepared and inoculated into 2-L glass bottles containing 1 L of medium, with six replicates for each salinity condition. The cultures were incubated at 25°C and 42°C, respectively. During the mid-exponential growth phase, when approximately 70 μM of the initial 100 μM ammonium had been oxidized, each culture was filtered through a 0.1-μm pore size mixed cellulose ester filter (Advantec, Japan). Filters were immediately flash-frozen in liquid nitrogen and stored at -80°C.

Total RNA was extracted using the AllPrep DNA/RNA Mini Kit (Qiagen, Germany) according to the manufacturer’s protocol. RNA integrity was assessed using an Agilent 2100 Expert Bioanalyzer (Agilent Technologies). cDNA libraries were constructed using the Nugen Universal Prokaryotic RNA-Seq Library Preparation Kit and sequenced on a NovaSeq6000 platform (Illumina) by LabGenomics Inc., Korea. Quality control of the sequencing reads was conducted using FastQC (v0.11.8) [67], and adaptor and quality trimming were performed using BBTools (v39.06) [86]. Reads mapping to rRNA sequences were removed using SortMeRNA (v2.1) [87]. The remaining reads were then aligned to the genomes of “*Ca.* N. chungbukensis” MY2 and *N. viennensis* EN76 using STAR (v2.7.11a) [88], and the mapped reads for each gene were counted using HTSeq (v2.0.5) [89]. Gene expression levels were calculated and presented as transcripts per kilobase million (TPM). Differential expression analysis was conducted using the DESeq2 package (v1.40.2) [90] in R (v4.3.2) [91]. *p*-values were calculated using a two-sided Wald test, and multiple comparisons were adjusted using the Benjamini-Hochberg method, as implemented in DESeq2.

Following protein extraction and enzymatic digestion with trypsin, peptide mixtures were analyzed using a nano-flow high-performance liquid chromatography (HPLC) system (UltiMate 3000 RSLCnano, Dionex, Thermo Fisher Scientific). Peptides were first loaded onto a C18 trap column (PepMap100, 300 μm × 2 cm, 5 μm particle size, nanoViper, Thermo Fisher Scientific), and subsequently separated on a C18-reverse-phase analytical column (Acclaim PepMap® 100, 75 μm × 25 cm, 3 μm particle size, nanoViper, Thermo Fisher Scientific). The separated peptides were introduced into a Q Exactive HF mass spectrometer (Thermo Fisher Scientific) coupled with a TriVersa NanoMate nano-electrospray ionization source (Advion, Harlow, UK) operating in LC chip coupling mode. Mass spectrometry acquisition parameters, including the LC gradient, ionization settings, and scan modes, were followed previously established protocols [92].

Although proteomic analysis of the two AOA strains under different salinity conditions enabled the detection of some proteins corresponding to highly expressed genes identified in the transcriptomic analysis, a more detailed interpretation was not pursued due to the limited depth of the obtained proteome data.

## Results and discussion

### Responses of soil ammonia oxidizers to low salinity

The effects of varying salinity levels (with ECs ranging from 60 to 780 μS/cm) on nitrification activity and the ammonia-oxidizing community were investigated using soil slurry microcosms derived from grassland and agricultural soils. The ECs of the soil slurry microcosms prepared with 0× MWM (without basic salts; EC ∼60 μS/cm) and 1× MWM (EC ∼120 μS/cm) were comparable to those observed in low-salinity soil environments, aligning with the 50th percentile of soil ECs (Supplementary Fig. S1; Table S1) [15]. For comparison, soil slurry microcosms prepared with 10× MWM had an EC value of ∼780 μS/cm, corresponding to the 90th percentile of soil ECs. Microcosms were further amended with or without ATU, a specific inhibitor of bacterial ammonia oxidation by AOB and CMX [3, 93], to differentiate the response of AOA from that of bacterial ammonia oxidizers. Activities of ammonia oxidizers in the microcosms were assessed through the measurement of nitrate production from nitrification and changes in the abundances of AOA, AOB, and CMX following ammonia oxidation (Fig. 1).

**Fig. 1.**
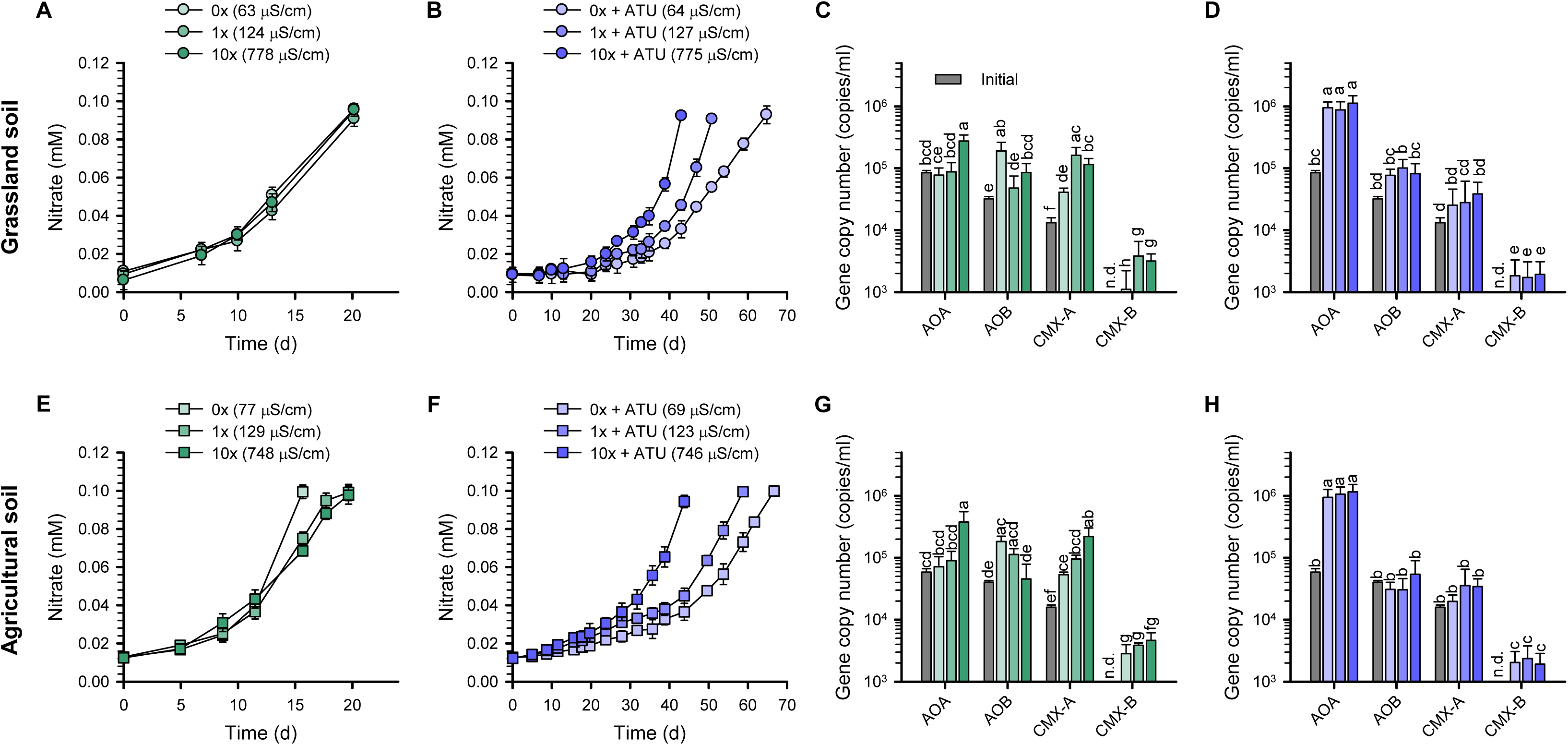
Changes in nitrate concentrations and the abundances of ammonia oxidizers during incubation of soil slurry microcosms. Results from microcosms using grassland soil (A–D) and agricultural soil (E–H) are shown. Nitrate concentrations (A, B, E, F) and the corresponding qPCR results (C, D, G, H) are presented. Microcosm samples were collected at the endpoint of ammonia oxidation (A, B, E, F) for downstream qPCR and 16S rRNA gene amplicon analysis. Panels B, F, D, and H show results from ATU-treated microcosms. Error bars represent standard deviations of biological replicates (n ≥ 3). Different letters denote statistically significant differences between treatments and the initial time point (two-way ANOVA followed by Tukey’s test, *p* < 0.05).

Nitrate production rates were comparable across all salinity levels in both soil microcosms during the 20-day incubation (Figs. 1A and E). However, salinity-dependent differences in the abundances of *amoA* genes affiliated with distinct ammonia oxidizer groups became evident after incubation. The abundance of the AOA *amoA* gene increased significantly only under the 10× MWM conditions, whereas that of CMX-A gradually increased with rising salinity (*p* < 0.05; Figs. 1C and G). In contrast, the abundance of the AOB *amoA* gene was highest under 0× MWM conditions in both soils, displaying a trend opposite to that of CMX-A and AOA (*p* < 0.05; Figs. 1C and G). The abundance of the CMX-B *amoA* gene remained a minor component, with no significant differences observed among salinity levels after incubation. Overall, while ammonia oxidation rates were comparable among salinity conditions (Figs. 1A and E), the growth responses of ammonia oxidizers varied: AOB was more abundant under lower salinity, whereas AOA and CMX-A became more prominent under higher salinity (Figs. 1C and G).

The response of AOA to salinity was further examined using ATU. Ammonia oxidation was delayed in the presence of ATU in both soil microcosms (Figs. 1B and F) compared to the control without ATU (Figs. 1A and E). AOA specific growth rates, calculated based on nitrate production [63], were lower under 0× and 1× MWM conditions (0.015 d^-1^ and 0.020 d^-1^, respectively, in both soils) compared to 10× MWM conditions (0.027 d^-1^ and 0.032 d^-1^ in grassland and agricultural soils, respectively), indicating that AOA growth is sensitive to lower salinity (Figs. 1B and F). This trend is consistent with the observed higher abundance of AOA at higher salinity in the absence of ATU (Figs. 1C and G). As expected, AOA were more abundant than AOB and CMX (*p* < 0.05; Figs. 1D and H) across all salinity conditions in both soils, likely due to the inhibitory effect of ATU on bacterial ammonia oxidation. Nevertheless, AOB and CMX-A *amoA* gene copies showed minor increases under ATU treatment (Figs. 1D and H), indicating that inhibition was not complete or may have weakened during the incubation period, possibly due to degradation or inactivation of ATU within the microcosm [94].

### Soil ammonia-oxidizing community analysis

To complement the qPCR results from the soil slurry microcosm experiments, 16S rRNA gene amplicon sequencing was conducted to analyze the ammonia-oxidizing community in samples collected initial and after the oxidation of added ammonia was complete (Fig. 2). In the absence of ATU, the relative abundance of the genus “*Ca.* Nitrosotenuis” within AOA-NpF significantly increased in response to elevated salinity in both soils (*p* < 0.05; Fig. 2A and C). This increase was primarily driven by ASV1979, which on average comprised 93% of the reads assigned to “Ca. Nitrosotenuis” in each sample and is identical in sequence to the corresponding 16S rRNA gene region of “*Ca.* N. chungbukensis” MY2 (Supplementary Fig. S2A; Dataset S2). The relative abundance of the genus *Nitrososphaera* within AOA-NsF in agricultural soil was also significantly higher under 10× MWM than under 0× MWM (Fig. 2C). In contrast, members of the *Nitrosomonas oligotropha* lineage within AOB were most abundant under lower salinity conditions, showing a significant response under those conditions in both soils (*p* < 0.05; Fig. 2A and C). ASV53, which is identical in sequence to *N. oligotropha* Nm45, was the dominant variant within this lineage, accounting for an average of 52% of the reads across samples (Supplementary Fig. S2B; Dataset S2). In agricultural soil, *Nitrosomonas communis* within AOB showed the highest relative abundance under 1× MWM, replacing the *N. oligotropha* lineage, which showed the highest relative abundance under 0× MWM (Fig. 2C). At 10× MWM, both lineages exhibited lower relative abundances than at their respective peaks under lower salinity conditions (Fig. 2A and C). Due to limitations in 16S rRNA gene resolution, CMX and nitrite-oxidizing bacteria within the genus *Nitrospira* are not discernible from each other [3, 4, 95] and were therefore excluded from this analysis. Most of the *Nitrospira* reads belonged to lineage II, which was consistently detected in both soil types at relative abundances ranging from 0.7% to 4.1%, regardless of the salinity (Supplementary Fig. S2C; Dataset S2). In the presence of ATU, the relative abundance of the genus “*Ca.* Nitrosotenuis” markedly increased in both soil microcosms following ammonia oxidation, dominating over other ammonia oxidizers regardless of salinity levels (Figs. 2B and D; *p* < 0.05), despite lower AOA specific growth rates under 0× and 1× MWM conditions compared to 10× MWM conditions (Figs. 1B and F). The same ASV1979, which was the predominant variant under ATU-free conditions, remained predominant in the presence of ATU (Supplementary Fig. S2A; Dataset S2).

**Fig. 2.**
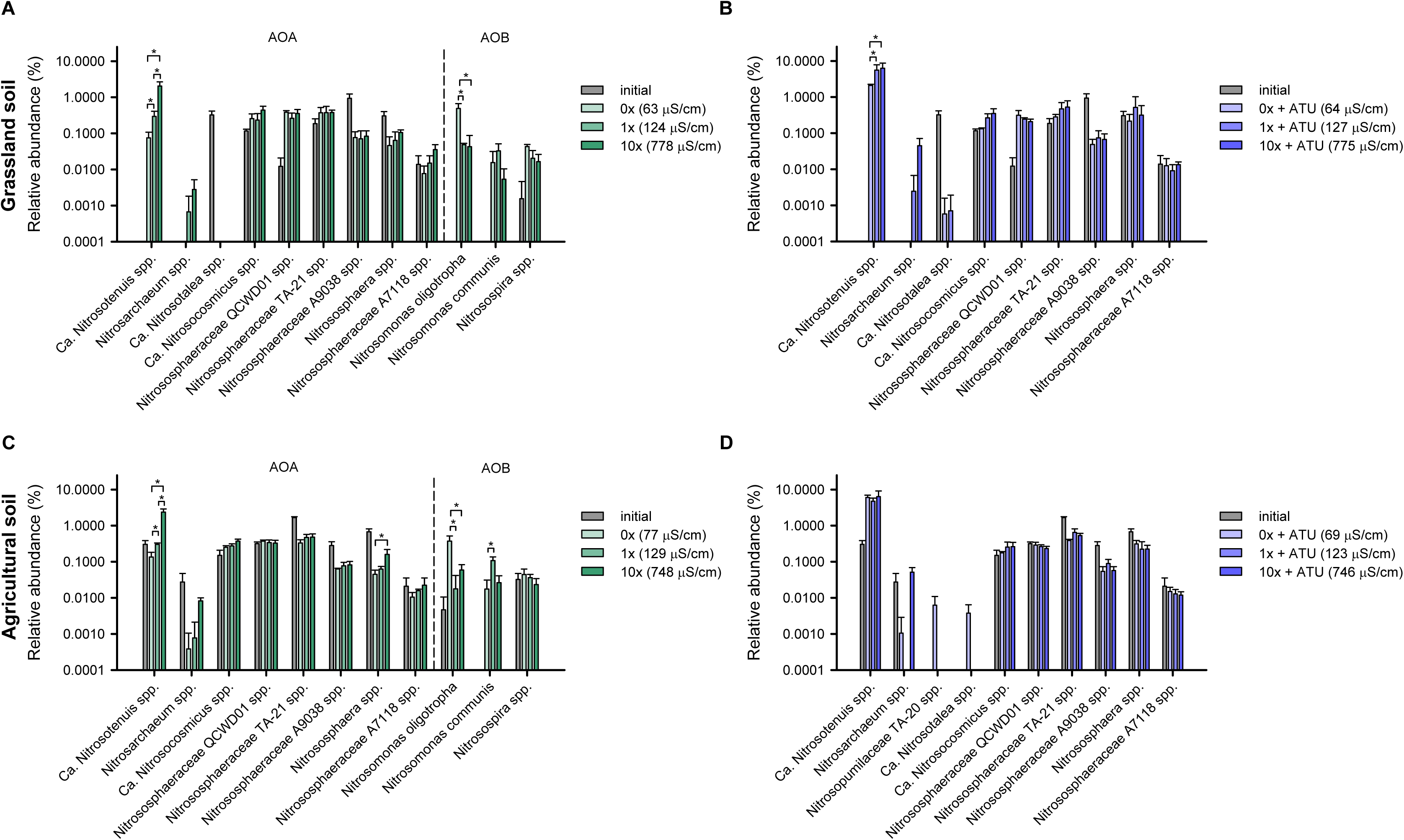
Relative abundances of ammonia oxidizers in soil slurry microcosm samples. The relative abundances (as % of total 16S rRNA gene reads) of ammonia oxidizers at the genus level are shown for microcosms derived from grassland soil (A, B) and agricultural soil (C, D). Panels B and D show results from the ATU-treated microcosms. Taxa detected in only a single replicate across all treatments or with abundances below 0.002% were excluded from the analysis. Error bars represent standard deviations of biological replicates (n ≥ 3). Initial samples were excluded from statistical analyses. Statistically significant differences between treatments for each ammonia oxidizer are indicated (two-way ANOVA, Tukey’s test, *p* < 0.05). Detailed relative abundances at the ASV level are available in Dataset S2.

### Responses of lake ammonia oxidizers to salinity change

To investigate the response of ammonia oxidizers to salinity changes in freshwater lakes, microcosms were prepared using unfiltered natural water samples from Lake Soyang and Lake Daecheong (Supplementary Table S2), each retaining its intrinsic salinity. The microcosms containing supplements such as trace metals but no added basic salts (0× MWM; Supplementary Table S1) exhibited ECs of ∼170 μS/cm, representing lower salinity conditions. In contrast, the addition of 10× MWM salts yielded ECs of ∼870 μS/cm, representing higher salinity. In both lake water microcosms, ammonia oxidation was completed within 10–25 days of incubation, and nitrite production rates (in these experiments, nitrite accumulated instead of nitrate) remained consistent across salinity levels (Figs. 3A and E). This nitrite accumulation may reflect rapid AOB activity, with limited subsequent oxidation by nitrite-oxidizing bacteria. After ammonia oxidation was completed, AOB *amoA* gene abundance exceeded that of other ammonia oxidizers regardless of salt supplementation (Figs. 3C and G). This consistent dominance suggests that lake-adapted AOB had a competitive advantage relative to other ammonia oxidizers.

**Fig. 3.**
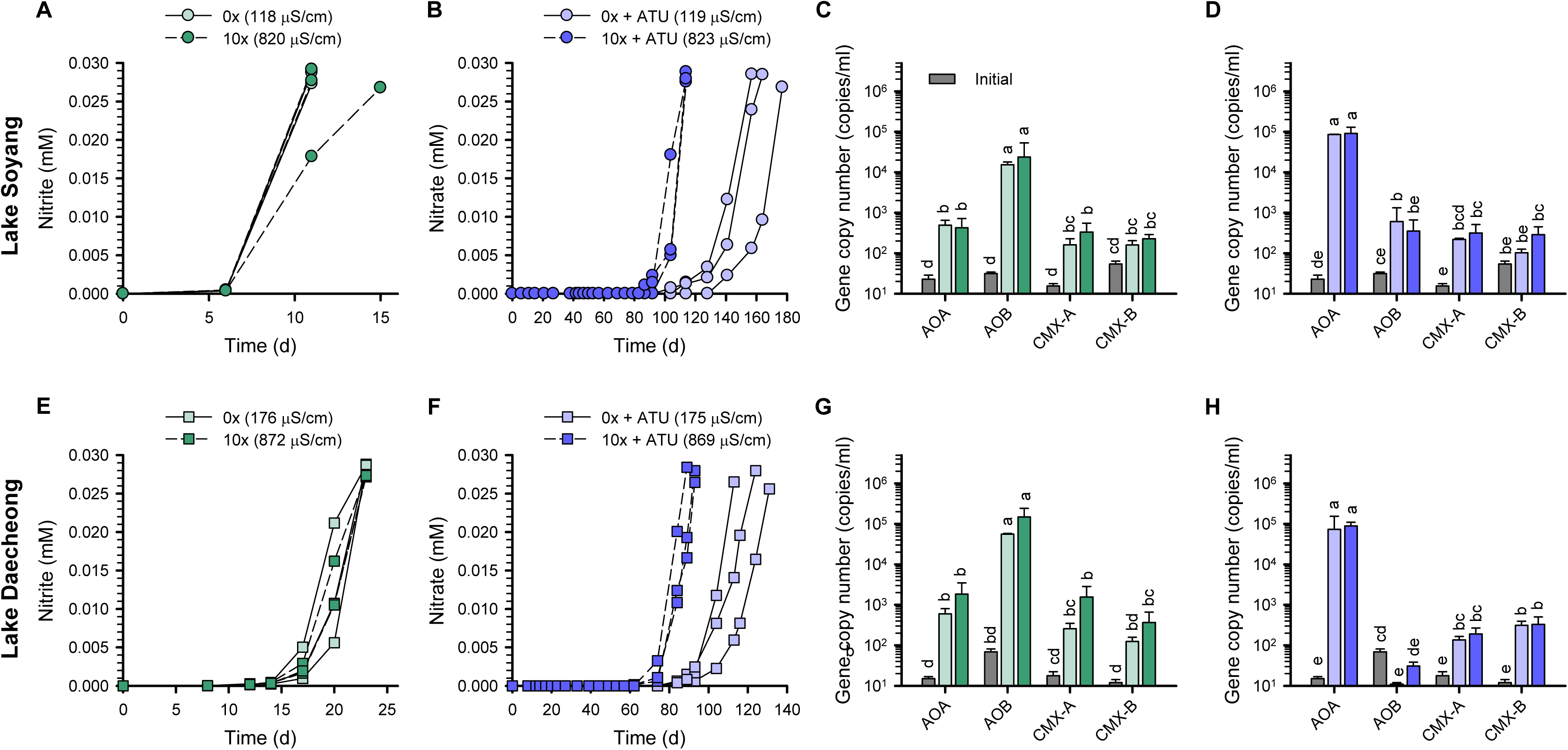
Changes in nitrite/nitrate concentrations and the abundances of ammonia oxidizers during incubation of lake microcosms. Results from microcosms using Lake Soyang (A–D) and Lake Daecheong (E–H), supplemented with 25 μM ammonia, respectively, are shown. In the absence of ATU, nitrite rather than nitrate accumulated (A, E). Each line represents an individual biological replicate, as high inter-replicate variability precluded meaningful averaging. Some lines overlap due to similar values (A, B, E, F). Microcosm samples were collected at the endpoint of ammonia oxidation (A, B, E, F) for downstream qPCR and 16S rRNA gene amplicon analysis. The corresponding qPCR results are shown in panels C, D, G, and H. Panels B, F, D, and H show results from ATU-treated microcosms. Error bars represent standard deviations of biological replicates (n ≥ 3). Different letters denote statistically significant differences between treatments and the initial time point (two-way ANOVA followed by Tukey’s test, *p* < 0.05).

In the presence of ATU, ammonia oxidation was delayed, requiring approximately 90 to 180 days for completion (Figs. 3B and F). Similar to patterns observed in soil microcosms, nitrate production was faster at elevated salinity (10× MWM). To determine whether lake water itself suppresses the nitrification activity of AOA, *Nitrosarchaeum koreense* MY1 was inoculated into microcosms containing unfiltered Lake Soyang waters with ATU (Supplementary Fig. S3). Upon inoculation of the strain, ammonia oxidation under 10× MWM conditions was completed much faster than in the uninoculated microcosms (Fig. 3B), within 26 days (Supplementary Fig. S3). However, ammonia oxidation remained minimal and incomplete under 0× MWM conditions. This demonstrates that AOA was not inhibited by lake water itself, while sensitivity to low salinity remained evident in the microcosms. Moreover, the prolonged delay in archaeal ammonia oxidation under high salinity conditions in the presence of ATU is likely attributable to the initially low viability of AOA cells or a long resuscitation time for dormant AOA cells in the original lake water. Although AOA *amoA* gene was predominant after ammonia oxidation regardless of salt supplementation (Figs. 3D and H), a slight increase in AOB and CMX-A/B *amoA* gene abundances was also observed under ATU treatment, similar to the soil microcosms (Figs. 1D and H), possibly reflecting incomplete inhibition or partial loss of ATU efficacy [94]. Together, these findings indicate that AOA in lake waters are subjected to stress under low salinity conditions, leading to reduced nitrification activity.

### Lake ammonia-oxidizing community analysis

Unlike in the soil microcosms, there was no significant difference in the relative abundance of ammonia oxidizers between low- and high-salinity conditions in lake microcosms in the absence of ATU. AOB affiliated with the *Nitrosomonas oligotropha* lineage were predominant in the ammonia-oxidizing community after the completion of ammonia oxidation in both lake microcosms (Figs. 4A and C). The composition of ASVs within the *N. oligotropha* lineage AOB was diverse across both lakes and salinity levels, with no single ASV dominating (Supplementary Dataset S2), suggesting extensive microdiversity at the ASV level, potentially reflecting their adaptation to oligotrophic and low salinity environments [96]. In contrast to the soil microcosms, which were dominated by the genus “*Ca.* Nitrosotenuis”, the genera *Nitrosarchaeum* (ASV271 and ASV909) and *Nitrosopumilus* (ASV135 and ASV827) within the AOA-NpF (Supplementary Fig. S2A; Dataset S2) were relatively abundant in lake microcosms (Figs. 4A and C), regardless of salinity. ASV271 and ASV909 were closely related (>98.9% identity) to *N. koreense* MY1, and ASV135 and ASV827 showed >99.5% identity to *Nitrosopumilus maritimus* SCM1 (Dataset S2). In the presence of ATU, and consistent with the soil microcosm results, AOA in the genus *Nitrosarchaeum* (ASV271 in Lake Soyang and ASV909 in Lake Daecheong) were dominant (Figs. 3B and F; 4B and D).

**Fig. 4.**
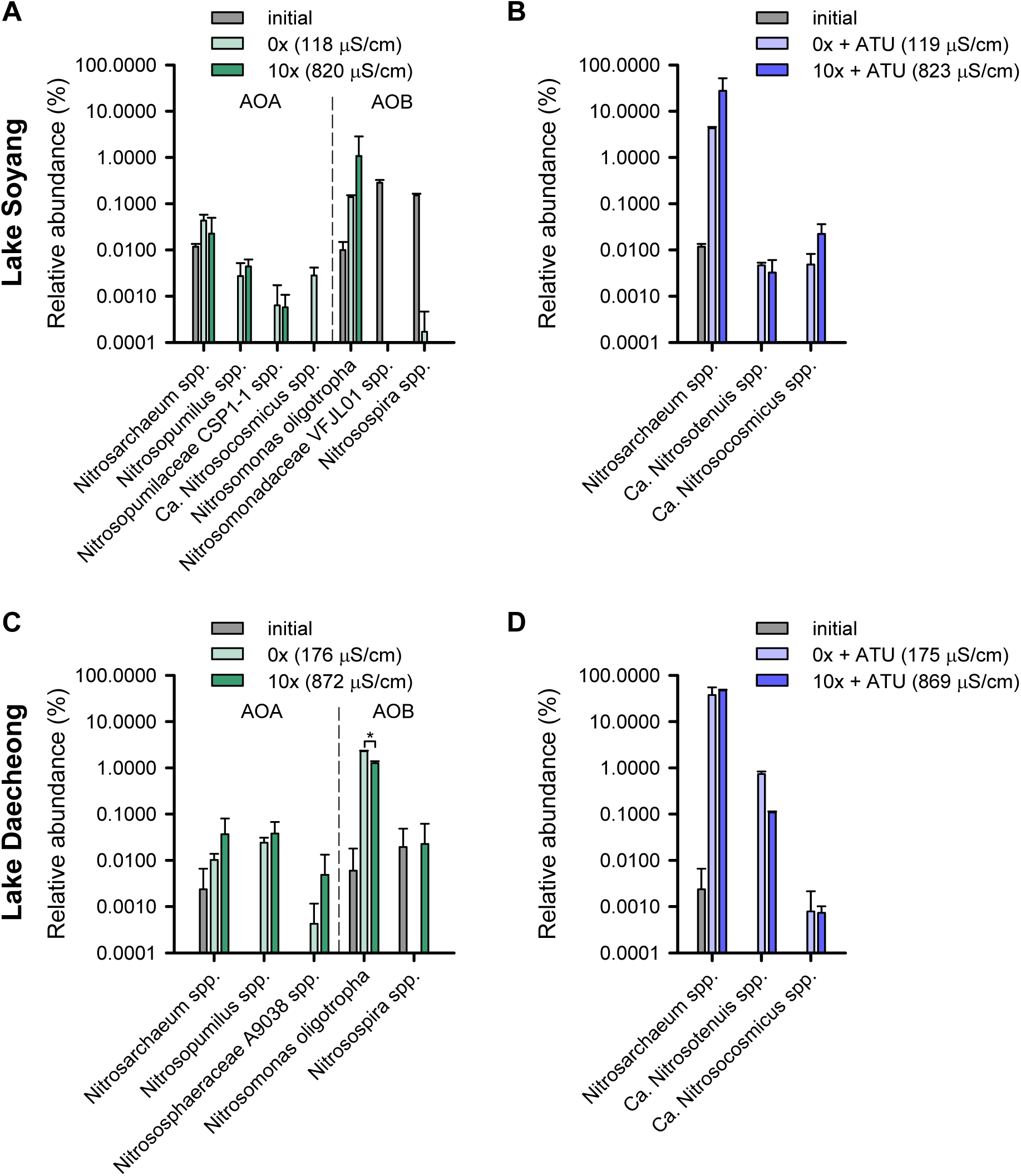
Relative abundances of ammonia oxidizers in lake microcosm samples. The relative abundances (as % of total 16S rRNA gene reads) of ammonia oxidizers at the genus level are shown for microcosms derived from Lake Soyang (A, B) and Lake Daecheong (C, D). Panels B and D show results from the ATU-treated microcosms. Taxa detected in only a single replicate across all treatments or with abundances below 0.002% were excluded from the analysis. Error bars represent standard deviations of biological replicates (n ≥ 3). Initial samples were excluded from statistical analyses. Statistical analysis was performed (two-way ANOVA followed by Tukey’s test, *p* < 0.05); however, no significant differences between treatments were observed for the taxa shown.

Taken together, results from the soil microcosms showed a significant increase in AOB abundance at lower salinity levels (Figs. 1C and G), possibly due to the inhibition of AOA or CMX-A (Figs. 1B, C, F, and G), thereby providing a competitive advantage to AOB. In contrast, AOA became the most dominant group after ammonia oxidation in soil microcosms under elevated salinity conditions (Figs. 1C and G). In the lake microcosms, AOB were predominant in both salinity levels, likely reflecting the low viability or delayed response of AOA and CMX (Fig. 3). The major AOB detected in both soil and lake microcosms were affiliated with the *N. oligotropha* lineage (Figs. 2A and C). The genera “*Ca.* Nitrosotenuis” and *Nitrosarchaeum* of the AOA-NpF were selectively responsive to high salinity in both soil and lake microcosms, respectively (Figs. 2, 4B and D).

### Effects of hypoosmolarity on the growth of ammonia oxidizers

The findings from microcosm experiments indicated that the nitrification activity of ammonia oxidizers is differentially affected by low salinity within the range typical of natural soil or lake environments (Supplementary Fig. S1). Thus, we hypothesized that chronic hypoosmolarity stress at low salinity levels affects AOA or CMX-A. To test this hypothesis, we used representative cultivated strains of AOA (AOA-NpF: *N. koreense* MY1 and “*Ca.* N. chungbukensis” MY2; AOA-NsF: *N. viennensis* EN76 and “*Ca.* N. oleophilus” MY3), AOB (*N. europaea* ATCC 19718), and CMX-A (“*Ca*. N. inopinata” ENR4). First, the growth of AOA-NpF strains (*N. koreense* MY1 and “*Ca.* N. chungbukensis” MY2) and AOA-NsF (*N. viennensis* EN76) was examined across a range of salinity levels, from 0× to 128× MWM (71–2,700 μS/cm) (Supplementary Fig. S4). The growth rates of the three AOA strains were generally lower at 0– 4× MWM (71–318 μS/cm) compared to 10× MWM, with statistically significant reductions observed for strains *N. koreense* MY1 and “*Ca.* N. chungbukensis” MY2, particularly at the lowest salinity levels. While *N. viennensis* EN76 also exhibited a significant reduction at 0× MWM, its growth rate declined less sharply across the salinity gradient, indicating a comparatively lower sensitivity to hypoosmolarity. At 10× MWM (795 μS/cm), the growth rates of “*Ca*. N. chungbukensis” MY2 and *N. viennensis* EN76 were comparable to those observed in AFM (3,540 μS/cm), a conventional medium for cultivating ammonia oxidizers [63]. Nonetheless, *N. koreensis* MY1 exhibited the highest difference between the two conditions. These results are consistent with the observed inhibition of ammonia oxidation by AOA under lower salinity levels (0× and 1× MWM) in both soil and lake microcosm experiments (see Figs. 1 and 3). Based on these findings, 1× MWM (128 μS/cm) and 10× MWM (795 μS/cm) were selected as hypoosmotic and control conditions, respectively, to assess the response of other ammonia oxidizers to hypoosmotic stress as they reflect lower and higher salinity levels in natural environments.

The inhibitory effect of low salinity conditions was more pronounced in AOA-NpF strains than in AOA-NsF strains. Specifically, at 1× MWM, the growth rates of AOA-NpF strains (*N. koreense* MY1 and “*Ca.* N. chungbukensis” MY2) were only 7.9% and 22.5%, respectively, of those observed under 10× MWM (Fig. 5). In contrast, AOA-NsF strains (*N. viennensis* EN76 and “*Ca.* N. oleophilus” MY3) exhibited less inhibition at 1× MWM compared to 10× MWM, retaining 82.1% and 76.4% of their growth rates, respectively (Fig. 5). These differences in inhibition across clades align with the findings from soil microcosm experiments, where the relative abundance of AOA-NpF greatly increased in a salinity-dependent manner (Figs. 2A and C). This pattern is further supported by slower growth rates of dominant AOA-NpF under lower salinity with ATU, as observed in both soil and lake microcosms (Figs. 1–4). Meanwhile, the growth of the AOB strain, *N. europaea* ATCC 19718, did not differ significantly between 1× MWM and 10× MWM, supporting observations from soil and lake microcosms in which AOB remained active under low-salinity conditions (63–176 μS/cm) (Figs. 1–4). Similarly, the growth of the CMX-A strain, “*Ca*. N. inopinata” ENR4, showed no significant difference between 1× MWM and 10× MWM (Fig. 5), which contrasts with the CMX-A growth inhibition under low-salinity soil microcosms (Figs. 1C and G). “*Ca*. N. inopinata” ENR4 was isolated from a deep oil exploration well [4], which may account for physiological differences between this CMX-A and those CMX-A-members in moderate soil and lake environments. Future studies could explore the growth response of CMX-A to low salinity conditions using strains isolated from such environments. Overall, our results clearly indicate that among the ammonia oxidizers, AOA are especially sensitive to hypoosmolarity stress.

**Fig. 5.**
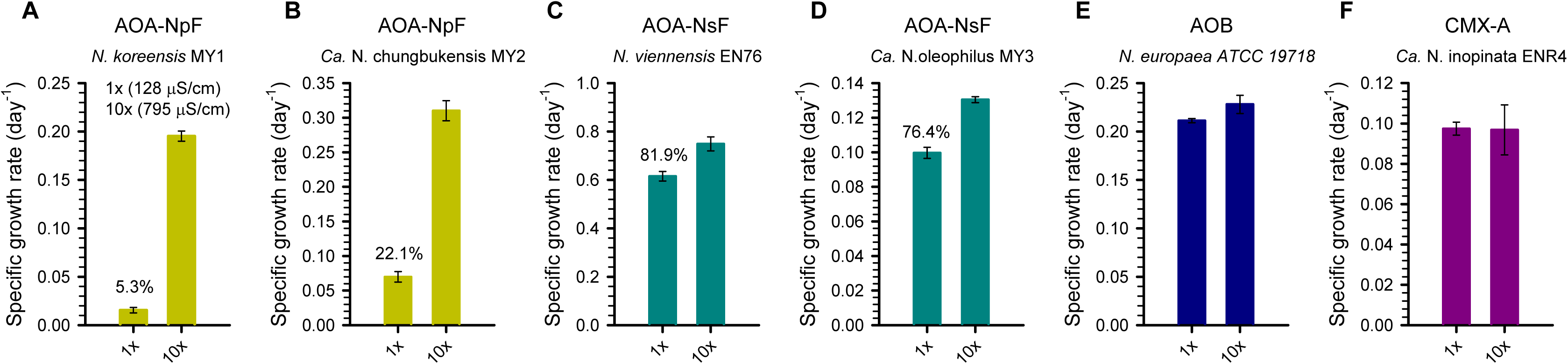
Growth of representative ammonia oxidizers under hypoosmotic conditions. Growth was compared between 1× and 10× MWM, representing the hypoosmotic and control conditions, respectively. The composition of MWM is provided in Table S1. Error bars represent the standard deviation of biological replicates (n ≥ 3). For strains exhibiting significantly reduced growth under hypoosmotic conditions, the percentage decrease in growth rate relative to the control is shown (two-tailed t-test*, p* < 0.05).

### Comparative genomic and gene expression analyses

Two key cellular properties may determine the sensitivity of microorganisms to hypoosmotic stress: 1) osmoregulatory systems that maintain ionic and osmotic balance [48, 49] and 2) cell surface structures that provide resistance to turgor pressure. Hypoosmotic shock imperatively leads to an influx of water across the cell membrane, increasing cell volume and turgor pressure. In the absence of a rigid cell wall, this pressure ultimately results in membrane rupture and loss of structural integrity [50]. To elucidate the mechanisms underlying the adaptation of ammonia oxidizers to hypoosmolarity, we compared the genomic repertoires of osmoregulatory systems across AOA-NpF, AOA-NsF, AOB, and CMX (Supplementary Fig. S5; Dataset S1) and performed transcriptomic and proteomic analyses. Compared to AOA, AOB, and CMX genomes contain various sensing and transport systems, primarily functioning in K^+^ or Na^+^ transport [51], which may provide greater flexibility in osmoregulation under hypoosmotic conditions (see Supplementary Text for details).

To reveal the mechanisms underlying AOA sensitivity to hypoosmotic stress, transcriptomic and proteomic responses of *N. viennensis* EN76 and “*Ca.* N. chungbukensis” MY2 cells grown at 1× and 10× MWM were comparatively analyzed. Out of 3,114 genes recovered from the transcriptome analysis of *N. viennensis* EN76, only 26 and 121 genes were significantly upregulated and downregulated at 1× MWM, respectively, based on a 1.5-fold change with an adjusted *p*-value < 0.01 (Supplementary Datasets S3 and S5). In the case of “*Ca.* N. chungbukensis” MY2, 139 and 168 genes were significantly upregulated and downregulated at 1× MWM, respectively, out of 2,019 genes recovered from the transcriptome analysis (Supplementary Datasets S4 and S5). Most central metabolism genes involved in ammonia oxidation, electron transport chain, CO2 fixation, TCA Cycle, and pentose phosphate pathway were constitutively expressed in both strains. However, genes differentially expressed under hypoosmotic conditions (1× MWM) belong to transport, cell surface, stress response, protein synthesis, proteolysis, biosynthesis, motility, and chemotaxis (Fig. 6; Supplementary Datasets S3–S5).

**Fig. 6.**
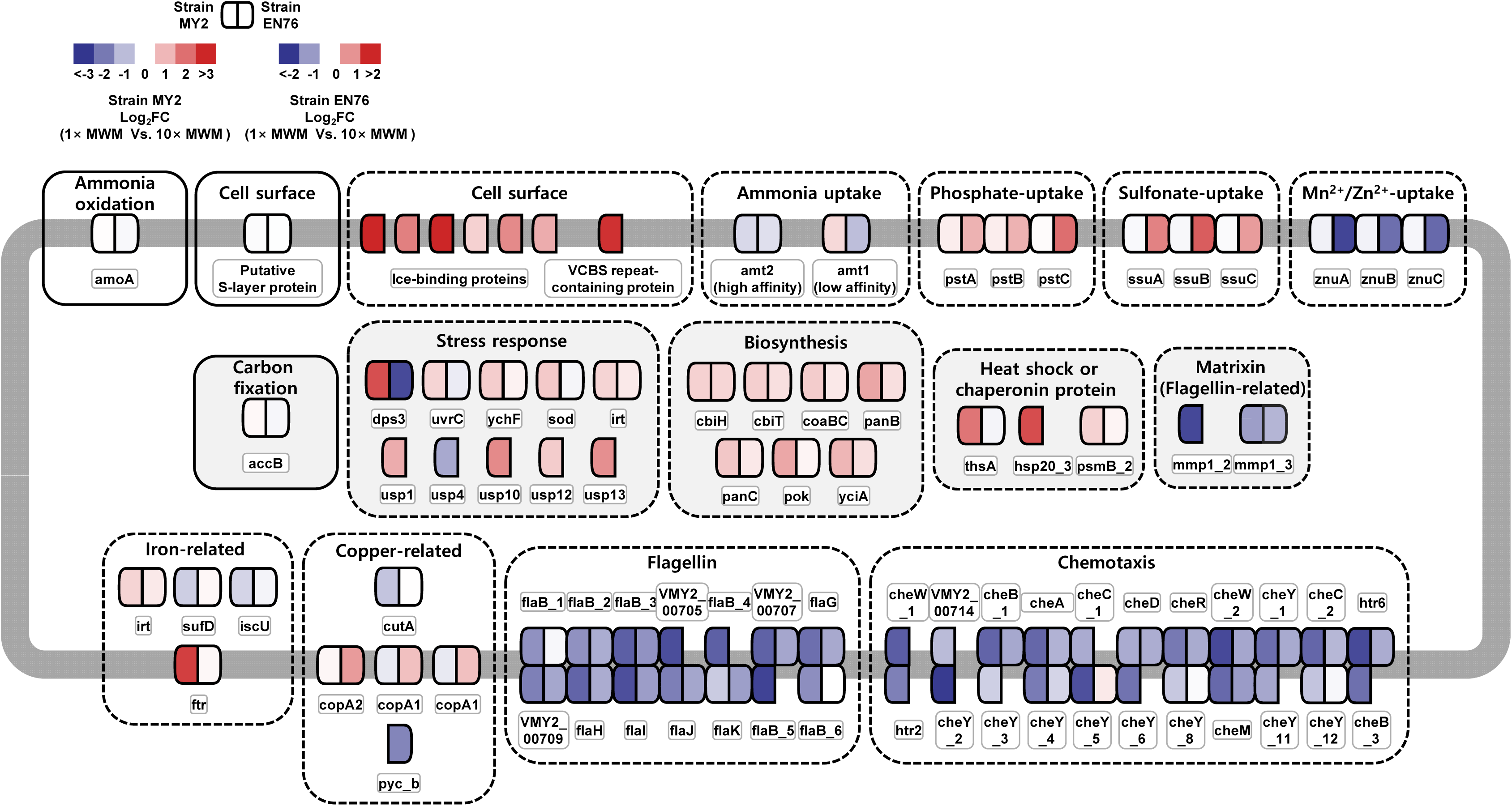
Differential gene expression under varying salinity conditions. A schematic overview of differentially expressed genes in “*Ca*. Nitrosotenuis chungbukensis” MY2 and *Nitrososphaera viennensis* EN76 under 1× MWM (128 μS/cm) vs. 10× MWM (795 μS/cm). Solid-bordered boxes denotes constitutively expressed genes; dashed-bordered boxes represent differentially expressed genes grouped by function. The color scale indicates higher gene expression in cells grown under 1× MWM (red) or 10× MWM (blue). The intensity of each color reflects the relative magnitude of expression change (log2fold change). See the Methods section for details on transcriptome sample preparation.

#### Transporters

In *N. viennensis* EN76, Genes encoding nutrient transporters were the most differentially expressed genes dependent on salinity. The nutrient transporter genes upregulated under 1× MWM were related to the transport of copper (*cop*), phosphate (*pst*), and aliphatic sulfonate (*ssu*) (Fig. 6; Supplementary Dataset S5). Conversely, genes for low-affinity ammonium transporter (*amt2*) and ABC-type Mn^2+^/Zn^2+^ transport system (*znu*) were downregulated under 1× MWM. In the case of “*Ca.* N. chungbukensis” MY2, high-affinity Fe^2+^/Pb^2+^ permease (*ftr*) were upregulated under 1× MWM (Fig. 6; Supplementary Dataset S5).

While the rationale for the upregulation of genes involved in ionic nutrient and metal uptake under hypoosmolarity conditions remains unclear, we suspect that this may be due to ionic gradient disruption across the membrane [97–99] or moderately low cation bioavailability at low salinity. Contrary to expectations, the genes for many other osmoregulatory systems, including Trk and Kdp potassium transporters, mechanosensitive channels, and aquaporin analyzed above (Supplementary SI Text; Fig. S5; Dataset S1), were constitutively expressed in both AOA strains (Supplementary Dataset S5). Together, these transcriptional responses suggest that low salinity alters the expression of genes related to nutrient uptake more prominently than those involved in osmoregulatory systems.

#### Cell surface structure

The sensitivity of AOA to hypoosmolarity may also be attributed to archaeal-specific features such as their cell wall and envelope structures. In contrast to the bacterial cells, which have semirigid cell walls and a high turgor pressure [100–102] due to peptidoglycan layers, archaea contain S-layer as their main cell envelope structure, which may compromise AOA cells’ structural resilience under low salinity conditions. For example, haloarchaeal cells are exposed to dilute environments, they lose the integrity of their S-layer lattice due to lack of divalent cations [103, 104]. Also, AOA cells are notably vulnerable to physical stress during centrifugation and filtration, while AOB cells are not [78, 105–107]. Comparatively, the number of genes related to membrane biosynthesis and cell envelope functionality is significantly higher in AOA-NsF than in AOA-NpF [108]. Further, cells in the AOA-NsF are coccoid while those of the AOA-NpF are rod-shaped. *Nitrosopumilus maritimus* SCM1 (AOA-NpF) has a hexagonal (p6) symmetry S-layer lattice [109], whereas *N. viennensis* EN76 (AOA-NsF) has a p3 symmetry S-layer lattice [79]. Together, possible differences in osmoregulatory systems and cell wall structures may explain the higher sensitivity of AOA, especially AOA-NpF clade, to hypoosmolarity. However, genes encoding the S-layer were not differentially expressed in both AOA strains (Fig. 6; Supplementary Dataset S5).

In contrast to *N. viennensis* EN76, which lacked differentially expressed genes related to cell surface proteins, most of the genes upregulated in “*Ca. N. chungbukensis*” MY2 cells grown under 1× MWM were associated with cell surface structures (Fig. 6; Supplementary Dataset S5). Notably, genes encoding ice-binding proteins (IBPs; also known as antifreeze proteins, AFPs), LPXTG-anchored collagen-like adhesins, and VCBS repeat-containing protein were highly expressed under this condition. Especially, five of the eight IBP genes ranked within the top 100 most highly expressed transcripts in 1× MWM, and their corresponding proteins alongside a LPXTG-anchored collagen-like adhesin were also detected in the proteome of 1× MWM-grown cells (Dataset S5). Despite growing cells of “*Ca.* N. chungbukensis” MY2 under 1× MWM at 30°C, the expression of IBP-like genes was unexpected. Four of the five IBP contain additional C-terminal domains of unknown function (Supplementary Fig. S6), which may indicate a broader role of these proteins, including “*Ca.* N. chungbukensis” MY2 cell envelope stabilization in dilute environments. In fact, IBP are suggested to be involved in protecting membrane stability [110–112]. Furthermore, the VCBS repeat-containing protein and LPXTG-anchored collagen-like adhesin may interact with the S-layer to form macromolecular surface assemblies [113, 114], which could serve as a protective mechanism to prevent structural damage in AOA cells under hypoosmotic stress. Therefore, their increased expression under low-salinity conditions may represent a physiological response to maintain cell envelope stability.

#### Others

Both strains downregulated the expression of motility-associated genes, including those encoding flagellin and chemotaxis-related proteins, which are generally clustered in the genome, under 1× MWM (14 and 22 genes, respectively) (Fig. 6; Supplementary Dataset S5). This downregulation has been widely reported under various stress conditions and is often interpreted as a metabolic shift that prioritizes stress tolerance over motility [115–117]. Differentially expressed genes involved in stress response and signal transduction, protein synthesis and degradation, and biosynthesis are described in the Supplementary SI Text. These suggest that low-salinity stress may trigger a broader physiological adjustment affecting cellular homeostasis.

Together, our findings demonstrate that AOA cells differ fundamentally from AOB cells in osmoprotection strategies due to differences in: 1) the osmoregulatory transport system and 2) cell surface structure [100–105]. These differences likely contribute to the heightened sensitivity of AOA, especially AOA-NpF, to hypoosmolarity stress, as observed in this study. In addition, representative isolates from two AOA clades used in this study exhibited distinct transcriptional responses to hypoosmotic stress, reflecting their different degrees of sensitivity. *N. viennensis* EN76 exhibited limited transcriptional changes (mainly related to nutrient uptake), while “*Ca.* N. chungbukensis” MY2, closely related to the highly responsive AOA-NpF abundant in higher salinity soil and lake microcosms, showed broad transcriptional changes, indicative of severe stress under hypoosmotic conditions. Notably, our results indicate that osmoprotection under hypoosmolarity is closely linked to cell wall rigidity, and the lack of a peptidoglycan-like cell wall in AOA likely plays a significant role in their heightened sensitivity to this stress.

### Ecological relevance of hypoosmolarity

Salinity is a key determinant of microbial community structure in various terrestrial and aquatic environments [12]. A freshwater microcosm study using a forested watershed stream showed that microbial diversity declined at lower salinity levels (<350 μS/cm) [43]. In the same study, the relative abundance of the phylum *Nitrososphaerota*, harboring AOA, began to increase only when the conductivity exceeded 350 μS/cm [43], highlighting AOA sensitivity to hypoosmolarity. The upper bound of this range is comparable to the biological effect concentration of 300 μS/cm (HC05; the 5th percentile hazardous concentration) reported for other aquatic organisms [44, 45], suggesting a potential biological threshold for hypoosmolarity at which many organisms, including AOA, may experience osmotic stress.

Consistent with these environmental observations, our microcosm and physiological studies strongly support that AOA, especially members of AOA-NpF, experienced growth inhibition under hypoosmotic conditions (<300 μS/cm), whereas AOB were largely unaffected (Figs. 1–4; Supplementary Fig. S4). In particular, the genus *Nitrosarchaeum* (members of the AOA-NpF), exhibited significantly delayed growth under hypoosmotic conditions in microcosms from Lake Soyang and Lake Daecheong (Figs. 3 and 4). These lakes, like many others in Korea, are exorheic with relatively short water residence times (<0.75 yr) and receive continuous freshwater inflow, resulting in persistently hypoosmotic conditions (<180 μS/cm) [118–120]. Given the heightened sensitivity of AOA-NpF to low salinity, their persistence in such environments is likely limited. In contrast, AOB, which are more resistant to hypoosmolarity, may be favored under these conditions, as reflected in their higher activity in both soil and lake microcosms (Figs. 1 and 3). This pattern is consistent with observations from other freshwater systems [121–123], where AOB consistently dominate over AOA even in environments where ammonium concentrations would typically favor AOA [121–123]. Similar to the lakes analyzed in this study, many of these freshwater systems commonly harbor *Nitrosarchaeum* as the dominant AOA lineage [121, 122]. For instance, in Flathead Lake (average conductivity of 150–172 μS/cm [124]), where maximum NH_4_ ^+^ concentrations reach ∼0.7 μM, the AOB genus *Nitrosospira* accounted for ∼2% of the prokaryotic community, while *Nitrosarchaeum* represents only ca. 0.1% [121]. Although *Nitrosarchaeum* has a higher affinity for NH3 + NH ^+^ (*Km(app)* = 0.56 μM [125]) compared to the AOB genus *Nitrosospira* (*Km(app)* = ∼150 μM [126]), AOB still remained dominant. Given that the ambient ammonium concentration in the lake is much closer to the *Km(app)* of *Nitrosarchaeum*, this disparity suggests that factors beyond substrate affinity, such as salinity, may influence the composition of the ammonia-oxidizing community. Similarly, in high-altitude lakes of the Sierra Nevada, where conductivities range from 1.8–94.2 μS/cm [127] and NH_4_ ^+^ concentrations between 2.8–72 nM, AOB consistently outnumbered AOA [123].

Similar patterns have been observed in some large lakes with extended water residence times (>2.6 yr), such as the Laurentian Great Lakes, which exhibit a conductivity range of 79–334 μS/cm [128–132]. Here, *Nitrosospira* consistently dominates over *Nitrosarchaeum* at nearly all sampling stations [122], where NH4^+^ concentrations generally remain below 300 nM—a condition expected to be more favorable to AOA than AOB. However, in other large lakes with long residence times, such as those in Central Asia and Europe—for instance, Lake Baikal (109–121 μS/cm) and the hypolimnion of Lake Constance (339 μS/cm) [133]—“*Ca.* N. limneticus” was the dominant ecotype of AOA [134], prevailing over other ammonia oxidizers. It has been suggested that the geographical isolation and long geological history of these lakes contributed to the adaptation of “*Ca.* N. limneticus” clade to local conditions. Notably, in aquifer systems, where *Nitrosarchaeum*-type AOA are detected in high abundance, potentially originating from external environmental inputs, AOB nonetheless drive the majority of ammonia oxidation activity [135]. Genomic changes suggested to have occurred during the transition of *Nitrosopumilus* species from marine to freshwater remain insufficient to explain the mechanisms of adaptation to hypoosmotic environments [136]. Thus, cultivation and detailed investigation of the osmoregulatory mechanisms of “*Ca*. N. limneticus” are necessary to clarify its functional role in nitrogen cycling and its ecological significance in hypoosmotic lakes.

### Conclusions

Although EC in terrestrial and aquatic environments is much lower than in general microbiological media, the responses of ammonia oxidizers to chronic hypoosmotic stress remain poorly understood. In this study, we show that AOA, particularly members of AOA-NpF, are highly sensitive to low-salinity conditions commonly encountered in soil and freshwater environments. In contrast, AOB displayed greater tolerance to low salinities and frequently dominated under such conditions. Genomic and transcriptomic analyses suggest that members of AOA-NpF possess limited osmoprotective capacity, possibly due to the lack of (i) robust osmoregulatory systems and (ii) of a rigid cell surface structure. With climate change intensifying the hydrological cycle, leading to more frequent and prolonged episodes of freshwater influx, low salinity in soils and lakes will increasingly become prevalent. This highlights the importance of understanding the impact of chronic hypoosmotic stress on terrestrial and aquatic nitrogen cycling. Hypoosmolarity-induced shifts in ammonia-oxidizing community composition could have important implications for nitrogen management practices and strategies to mitigate N2O emissions in terrestrial and aquatic environments. This study highlights osmotic stress as a potentially underrecognized but ecologically relevant factor influencing ammonia-oxidizing community dynamics in low-salinity environments.

## Data Availability

The transcriptomic and 16S rRNA gene amplicon sequencing data have been deposited in the NCBI BioProject under the accession number PRJNA1265630. The mass spectrometry-based proteomics data have been deposited to the ProteomeXchange Consortium via the PRIDE [137] partner repository under the dataset identifier PXD064500.

## Acknowledgments

This work was supported by the NRF (National Research Foundation of Korea) grant funded by the Korean government (Ministry of Science and ICT) (2021R1A2C3004015) and Basic Science Research Program through NRF funded by the Ministry of Education (2020R1A6A1A06046235). J-HG was supported by the NRF grant funded by the Korean government (Ministry of Science and ICT) (RS-2023-00213601). MW was supported by the Austrian Science Fund (FWF) Cluster of Excellence Microbiomes drive Planetary Health (10.55776/COE7).

## Author contributions

J.-H.G., A.O., and S.-K.R. designed research. J.-H.G. and A.O. performed research. J.-H.G., A.O., and S.-K.R. analyzed data. J.-H.G., A.O., and S.-K.R. wrote the manuscript with contributions and comments from all co-authors.

## Competing interests

The authors declare no competing interests.

## Notes

### Competing Interest Statement

The authors have declared no competing interest.

