## Supplementary information for "Hypoosmolarity inhibits archaeal ammonia oxidation"

### Supplementary material for Hypoosmolarity inhibits archaeal ammonia oxidation Short title: Hypoosmolarity stress on archaeal nitrification

Joo-Han Gwak^a,1^, Adebisi Olabisi^a,1^, Ui-Ju Lee^a^, Christiana Abiola^a^, Seongjun Lee^a^, Hackwon Do^b^, Yun Ji Choi^c^, Jay-Jung Lee^d^, Man-Young Jung^c,e^, Nico Jehmlich^f^, Martin von Bergen^f,g^, Michael Wagner^h,i^, Samuel Imisi Awala^j^, Zhe-Xue Quan^k^, and Sung-Keun Rhee^a,*^

^a^Department of Biological Sciences and Biotechnology, Chungbuk National University, 28644 Cheongju, Republic of Korea ^b^Division of Life Sciences, Korea Polar Research Institute, 21990 Incheon, Republic of Korea
^c^Interdisciplinary Graduate Program in Advanced Convergence Technology and Science, Jeju National University, 63243 Jeju, Republic of Korea
^d^Geum River Environment Research Center, National Institute of Environmental Research, 29027 Okcheon, Republic of Korea
^e^Department of Biology Education, Jeju National University, 63243 Jeju, Republic of Korea
^f^Department of Molecular Systems Biology, Helmholtz Centre for Environmental Research–Zentrum für Umweltforschung GmbH, 04318 Leipzig, Germany
^g^Institute of Biochemistry, Faculty of Biosciences, Pharmacy and Psychology, University of Leipzig, 04103 Leipzig, Germany
^h^Centre for Microbiology and Environmental Systems Science, Department of Microbiology and Ecosystem Science, University of Vienna, Vienna, Austria
^i^Department of Chemistry and Bioscience, Aalborg University, Aalborg, Denmark
^j^Department of Biological Sciences, University of Calgary, Calgary, AB T2N 1N4, Canada
^k^School of Life Sciences, Fudan University, Shanghai, China

^1^J-H.G. and A.O. contributed equally to this work.

*Sung-Keun Rhee

**Keywords:** Hypoosmolarity, Osmotic stress, Nitrification, Freshwater ecosystems

**This PDF file includes:**

Supplementary Text

Supplementary Tables S1 to S3

Supplementary Figures S1 to S5

Legends for Supplementary Datasets S1 to S5

Supplementary References

**Other supplementary materials for this manuscript include the following:**

Datasets S1 to S5

### Supplementary Text

#### *Comparative genomic analysis of osmoregulation systems in AOM*

A distinct difference in K^+^ transport systems was observed between AOB and other ammonia oxidizers (AOA and CMX). Genes encoding K^+^ transport proteins—such as putative flavoprotein involved in K^+^ transport, K^+^ uptake Trk/Ktr system TrkAH (TC 2.A.38.1, 2.A.38.4), multicomponent K^+^:H^+^ antiporter Pha (TC 2.A.63.1), and K^+^:H^+^ antiporter Kef (TC 2.A.37.1)—were found exclusively in AOB (Dataset S1). Most bacterial ammonia oxidizers (AOB and CMX), in contrast to AOA, also encoded Na^+^:2H^+^ antiporter NhaA (TC 2.A.33) or Na^+^:H^+^ antiporter NhaC (TC 2.A.35), and Mg^2+^ transporter MgtE (TC 1.A.26.1.6) (Supplementary Fig. S5; Dataset S1). These systems primarily function in K^+^ or Na^+^ efflux and may provide greater flexibility in osmoregulation under hypoosmotic conditions [1], particularly for AOB and CMX.

The Trk system typically operates as a complex of TrkA and TrkH, where TrkA serves as a regulatory subunit [2]. In AOA, however, only *trkH* homologs were detected (Supplementary Fig. S5; Dataset S1). In the absence of TrkA, TrkH is likely to function as a non-selective cation channel permeable to K^+^ or Na^+^ [2, 3]. This configuration may lead to uncontrolled K^+^ flux, which could impair homeostatic control under hypoosmotic conditions. By contrast, the high-affinity K^+^ transporter KdpABC (TC 3.A.3.7; *K*_m_ = ~2 μM [4]) was identified in several genomes of AOA-NsF, AOB, and CMX (Supplementary Fig. S5; Dataset S1), including strains that exhibited little or no growth inhibition under low-salinity conditions (Fig. 5). The presence of this high-affinity K^+^ uptake system may provide a selective advantage in mitigating hypoosmotic stress.

The low-salinity-insensitive strain *N. europaea* ATCC 19718 (Fig. 5) encodes only the TrkAH system for K^+^ uptake, along with two K^+^:H^+^ antiporters (YbaL and Pha), a voltage-gated K^+^ channel, and the mechanosensitive channel MscK, which requires external K^+^ for activation (Supplementary Fig. S5; Dataset S1). K^+^ transport in *N. europaea* is electrogenically driven [5]. Notably, even after potassium depletion via diethanolamine treatment, respiration with NH₄^+^ retained 83% of the untreated activity, and the proton-motive force was preserved in the absence of added K^+^ [5]. These observations may indicate that strain *N. europaea* ATCC 19718 can sustain osmotic and energetic homeostasis with relatively low sensitivity to external K^+^ availability. The two K^+^:H^+^ antiporters could potentially support low-K^+^ adaptation by facilitating proton-coupled K^+^ uptake [5], particularly at concentrations above 50 μM (1× MWM; Fig. 5).

#### *Transcriptomic and proteomic responses of AOA to hypoosmolarity*

**Stress response and signal transduction:** The transcriptomes of “*Ca.* N. chungbukensis” MY2 displayed a significant number of upregulated genes involved in stress response and signal transduction in 1× MWM in contrast to those of *N. viennensis* EN76 (Fig. 6; Supplementary Dataset S5). Five genes of universal stress protein, four genes of heat shock protein, *uvrC* (nucleotide excision repair gene), *dps* (DNA protection against starvation, superoxide dismutase, and Fe^2+^-trafficking protein for iron homeostasis), and *ychF* (redox-regulated ATPase) were upregulated under 1× MWM. Several signal transduction protein genes were differentially expressed, showing both up- and downregulation.

**Protein synthesis and degradation**: In *N. viennensis* EN76, a gene for trypsin-like peptidase domain-containing protein was upregulated under 1× MWM. “*Ca*. N. chungbukensis” MY2 showed upregulation of genes functioning as chaperone, translocase, and elongation factor (Supplementary Dataset S5).

**Amino acid and cofactor biosynthesis**: Upregulation of genes involved in amino acids and NAD biosynthesis, such as 2-isopropylmalate synthase and quinolinate synthase, was observed under the 1× MWM in *N. viennensis* EN76. In contrast, many genes related to nucleotide, lipid, sugar, and vitamin(coenzyme) metabolism were upregulated in “*Ca*. N. chungbukensis” MY2 (Fig. 6; Supplementary Dataset S5).

### Supplementary Tables

#### Table S1. Composition of the mineral water medium (MWM)^a^ and artificial freshwater medium (AFM) used in this study.

| Medium | 0× | 0.5× | 1× | 2× | 4× | 8× | 10× | 16× | 32× | 64× | 128× | AFM |
| --- | --- | --- | --- | --- | --- | --- | --- | --- | --- | --- | --- | --- |
| Conductivity^b^ (μS/cm) | 71 | 96 | 128 | 188 | 318 | 629 | 795 | 900 | 1,169 | 1,707 | 2,700 | 3,540 |
| Basic salts |  |  |  |  |  |  |  |  |  |  |  |  |
| NaCl (mM) | - | - | - | 0 | 0 | 0.7 | 0.9 | 1.5 | 3.1 | 6.3 | 12.7 | 17.1 |
| KCl (mM) | - | - | - | 0.05 | 0.15 | 0.35 | 0.45 | 0.75 | 1.55 | 3.15 | 6.35 | 6.71 |
| MgSO_4_·7H_2_O (mM) | - | 0.022 | 0.045 | 0.09 | 0.19 | 0.4 | 0.5 | 0.5 | 0.5 | 0.5 | 0.5 | 1.97 |
| CaCl_2_·2H_2_O (mM) | - | 0.09 | 0.18 | 0.38 | 0.78 | 1.6 | 2 | 2 | 2 | 2 | 2 | 0.68 |
| Supplementary amendments |  |  |  |  |  |  |  |  |  |  |  |  |
| NH_4_Cl (mM) | 0.025–0.1 (for microcosms) or 0.025–1 (for isolates) | | | | | | | | | | | |
| KH_2_PO_4_ (mM) | 0.05 (for microcosms or isolates) or 1.47 for AFM | | | | | | | | | | | |
| NaHCO_3_ (mM) | 0.1 (for microcosms or isolates) or 2 for AFM | | | | | | | | | | | |
| HEPES (mM) | 1 (for isolates) or 2 (for microcosms) | | | | | | | | | | | |
| Pyruvic acid (mM) | 0.1 (for MY1, MY2, EN76 strains) or none (for the other strains) | | | | | | | | | | | |
| Trace elements^c^ (×) | 1 | | | | | | | | | | | |
| Ferric sodium EDTA (μM) | 7.5 | | | | | | | | | | | |
| Allylthiourea ATU (μM) | 50 (for microcosms) | | | | | | | | | | | |
| Total metal ions at pH 7.0 |  |  |  |  |  |  |  |  |  |  |  |  |
| Na^+^ (mM) | 0.35 | 0.6 | 0.6 | 0.6 | 0.6 | 1.3 | 1.5 | 2.1 | 3.4 | 6.9 | 10.4 | 21 |
| K^+^ (mM) | 0.01 | 0.01 | 0.05 | 0.1 | 0.2 | 0.4 | 0.5 | 0.8 | 1.36 | 3.2 | 4.51 | 8.18 |
| Mg^2+^ (mM) | 0 | 0.022 | 0.045 | 0.95 | 0.19 | 0.4 | 0.5 | 0.5 | 0.5 | 0.5 | 0.5 | 1.97 |
| Ca^2+^ (mM) | 0 | 0.09 | 0.18 | 0.38 | 0.78 | 1.6 | 2 | 2 | 2 | 2 | 2 | 0.68 |

^a^For the soil slurry microcosms and cultivation of isolates, MWM was prepared using distilled water. However, for the lake microcosms, MWM was prepared directly using lake water.

^b^at pH 7.0 with 1 mM NH4Cl, 0.05 mM KH2PO4, 0.1 mM NaHCO3, and 1 mM HEPES

^c^Trace elements were prepared as described by Widdel and Bak [6]

#### Table S2. Description of the location and properties of soil and lake samples for the microcosm.

| Sample | Grassland soil | Agricultural soil | Lake Soyang | Lake Daecheong |
| --- | --- | --- | --- | --- |
| Location (GPS) | 36°37‘36.81“ N,  127°27‘20.31“ E | 36°37‘26.76“ N,  127°27‘14.40“ E | 37°56‘56.8“ N,  127°49‘04.3“ E | 36°28‘26.40“ N,  127°29‘13.56“ E |
| Sampling date | 2023-11 | 2023-11 | 2024-03 | 2024-05 |
| Sampling depth (m) | 0.5 | 0.5 | 20 | 20 |
| Conductivity (μS/cm) | 24.6 | 120.9 | 81.2 | 147.5 |
| Residence time (yr) | - | - | 0.75 | 0.5 |
| Average volume (km^3^) | - | - | 2.9 | 0.79 |
| Temperature (°C) | - | - | 6 | 15 |
| pH | 5.78 | 7.94 | 6.98 | 6.70 |
| PO_4_^3-^ (mg·N·kg^-1^ soil or μM) | n.d. | n.d. | < 1 uM | < 1 uM |
| NO_2_^-^ (mg·N·kg^-1^ soil or μM) | n.d. | n.d. | n.d. | n.d. |
| NO_3_^-^ (mg·N·kg-1 soil or μM) | 9.8 | 13.6 | 94.2 | 91.9 |
| NH_4_^+^ (mg·N·kg-1 soil or μM) | 3.2 | 9.6 | 5.7 | 7.1 |

#### Table S3. Primers used for qPCR quantification of AOA, AOB, and CMX *amoA* genes.

| Target | Primer name | Sequence | Condition | Reference |
| --- | --- | --- | --- | --- |
| AOB amoA gene | amoA-1F | GGGGTTTCTACTGGTGGT | 95°C for 5 min; 40 cycles of 95°C for 30 s, 60°C for 30 s, 72°C for 30 s; 72°C for 5 min | [7] |
|  | amoA-2R | CCCCTCKGSAAAGCCTTCTTC |  |  |
| AOA amoA gene | CamoA-19f | ATGGTCTGGYTWAGACG | 95°C for 5 min; 40 cycles of 95°C for 30 s, 60°C for 30 s, 72°C for 30 s; 72°C for 5 min | [8] |
|  | CamoA-616r | GCCATCCABCKRTANGTCCA |  |  |
| CMX clade A amoA gene | cmxA_378F | GTGGTGGTGGTCBAAYTA | 95°C for 5 min; 40 cycles of 95°C for 30 s, 55°C for 30 s, 72°C for 30 s; 72°C for 5 min | [9] |
| CMX clade B amoA gene | cmxB_378F | GTACTGGTGGGCBAAYTT |  |  |
| CMX clade A and B amoA gene | cmxAB574r | GAAGCCCATRTARTCNGCC |  |  |

### Supplementary Figures

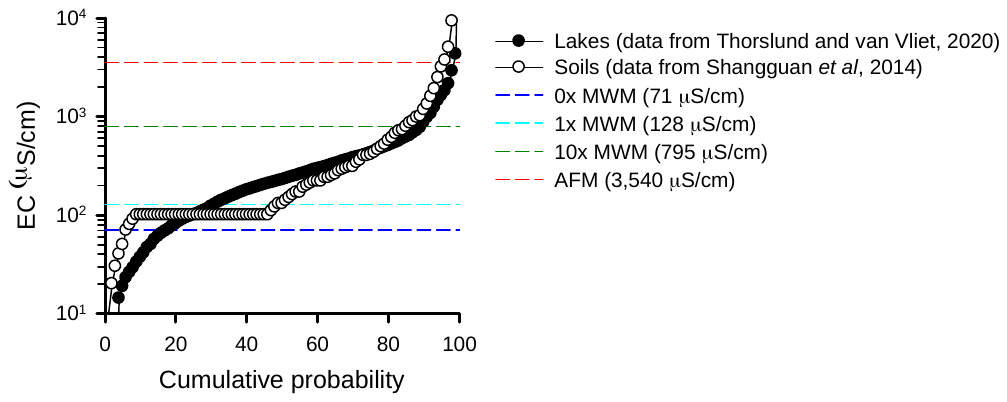

#### Fig. S1. Quantile function plot showing the electrical conductivity of lakes and soils on a global scale. Percentiles of electrical conductivity were calculated using the observed median values from lakes/reservoirs datasets (filtered for non-coastal locations) [10] and from soil datasets (filtered for non-zero values) [11] for lakes and soils, respectively. Dashed lines indicate the electrical conductivity of the media used in this study. Details on the composition of MWM and AFW are provided in Table S1.

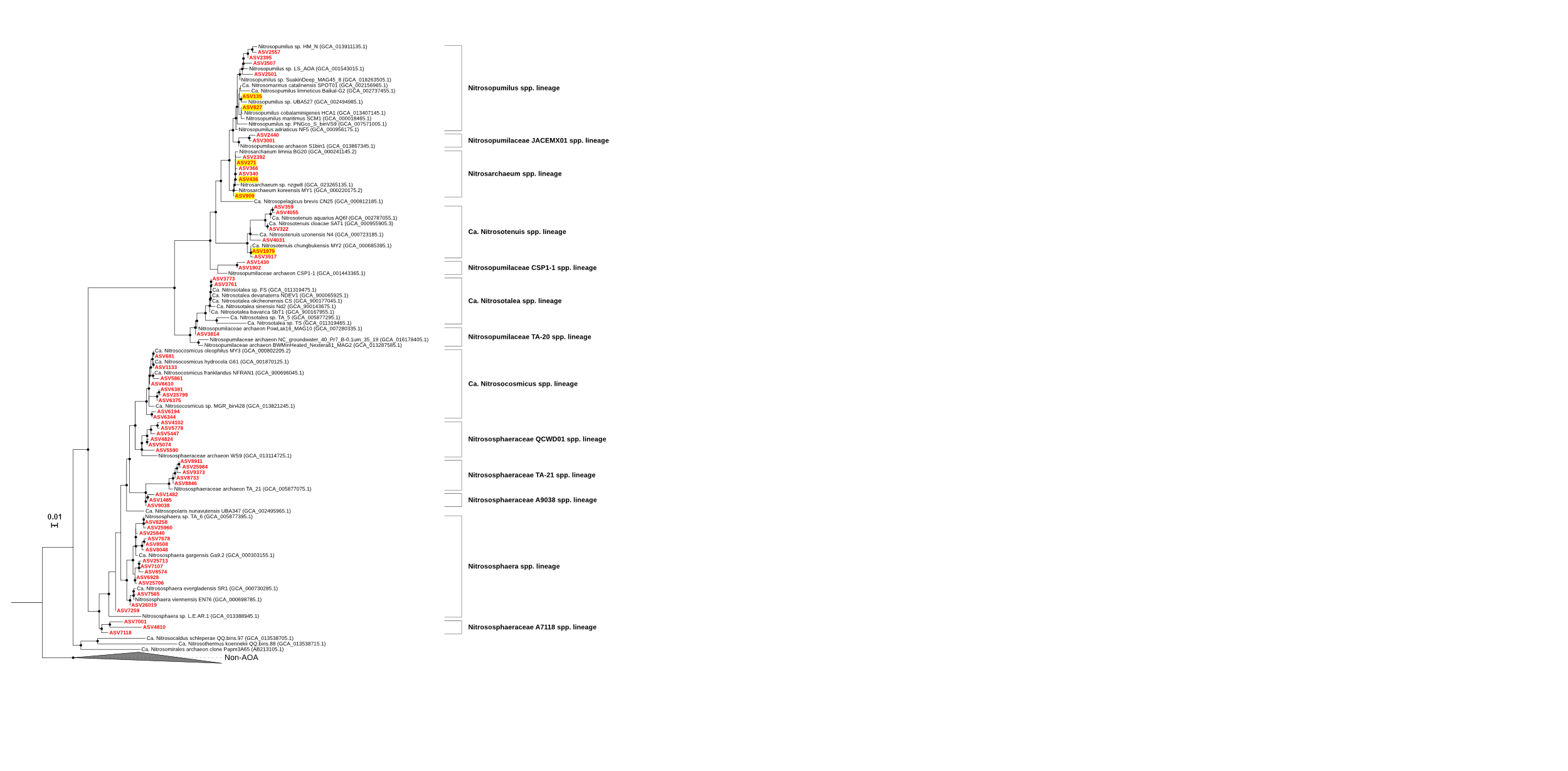

**A**

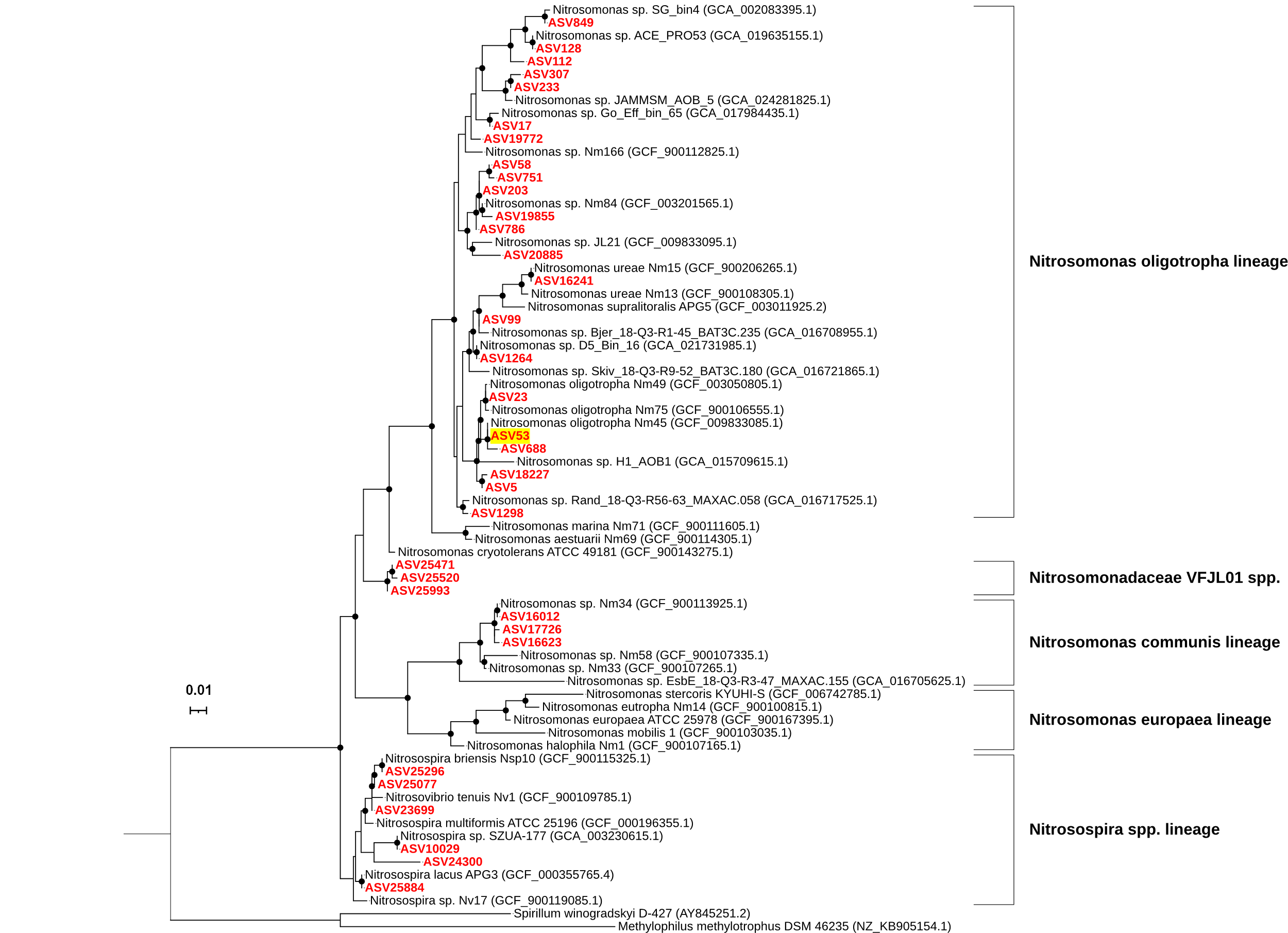

**B**

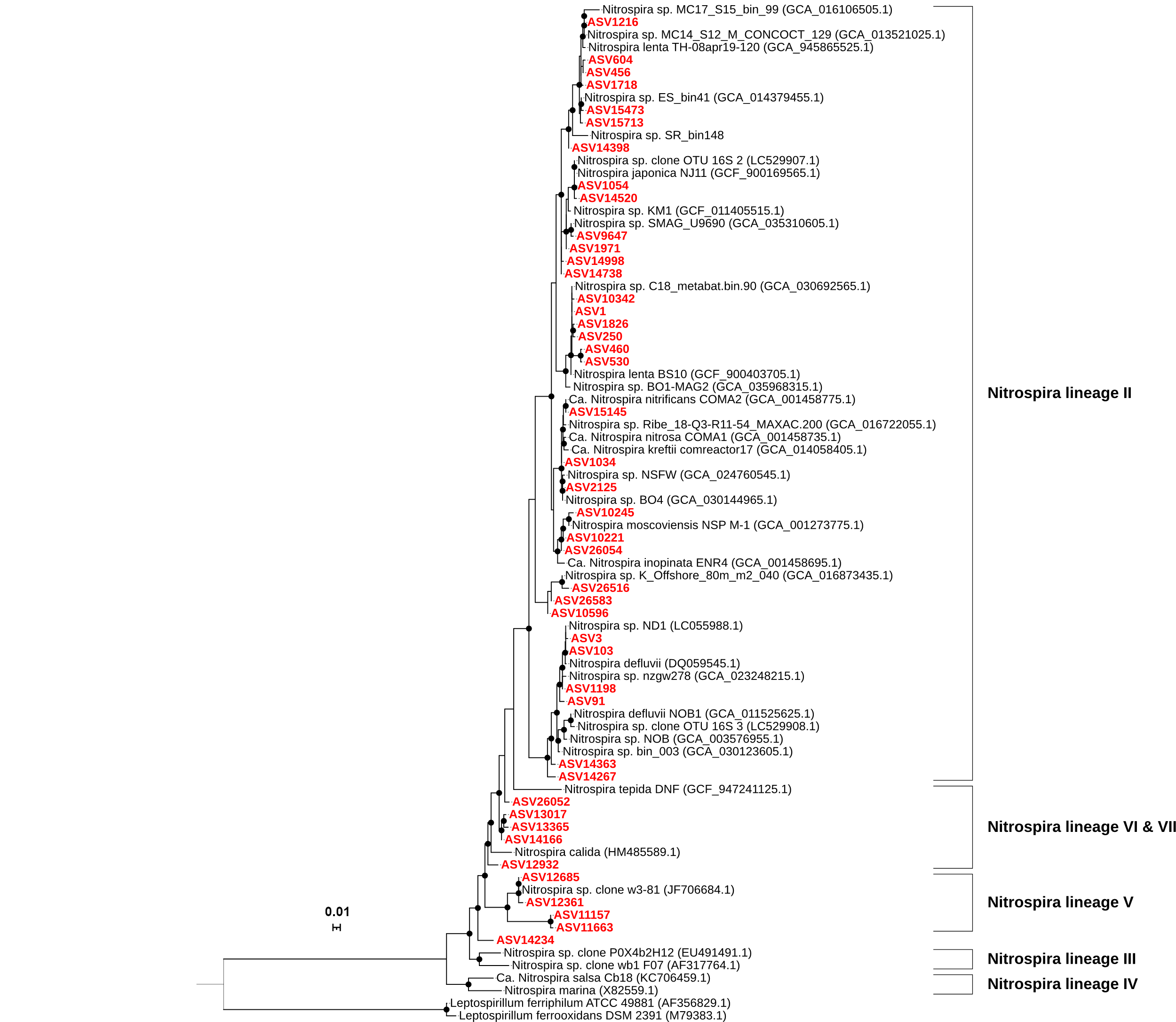

**C**

#### Fig. S2. 16S rRNA gene-based phylogenetic trees of ammonia oxidizers and the genus *Nitrospira*. Phylogenetic trees were constructed based on 16S rRNA gene sequences corresponding to the ASVs shown in Figures 2 and 4. (A) Ammonia-oxidizing archaea (AOA), (B) Ammonia-oxidizing bacteria (AOB), (C) Members of the genus *Nitrospira*. Black circles indicate nodes with ≥70% bootstrap support. ASV sequences were aligned, and phylogenetic trees were inferred using the methods described in the Materials and Methods section. ASVs mentioned in the main text are highlighted in yellow. The models SYM+I+R4 (A), TPM3+I+R3 (B), and TIM3+F+I+R3 (C) were selected by ModelFinder [12] and used for tree inference.

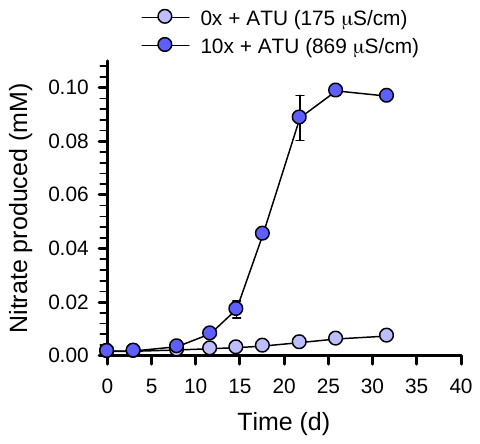

#### Fig. S3. Nitrate accumulation by *Nitrosarchaeum* *koreense* MY1 in Lake Soyang microcosms. *N. koreensis* MY1 (~10^7^ cells ml^−1^) was inoculated into unfiltered Lake Soyang water under different salinity conditions: 0× (MWM without basic salts) and 10× MWM. ATU was added to inhibit bacterial ammonia oxidation. Nitrate accumulation was monitored over time. Under 10× MWM, nitrification proceeded efficiently, with complete ammonia oxidation within 26 days, whereas a significant delay was observed under 0× MWM. Error bars indicate standard deviations of biological replicates (n ≥ 3).

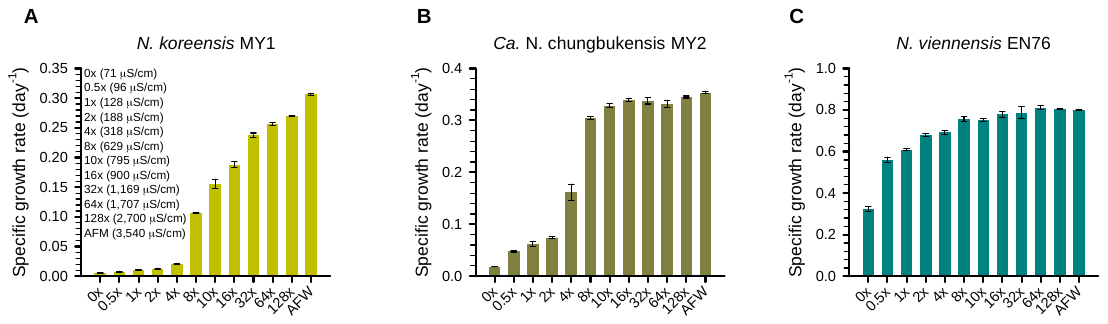

#### Fig. S4. Growth of ammonia oxidizing archaea strains in various salinity conditions. The composition of MWM (0–128×) and AFW are detailed in Table S1. Error bars represent the standard deviation for n ≥ 3 biological replicates. Significant differences between treatments for each nitrifier are indicated by different letters (one-way ANOVA, Tukey’s test, *p* < 0.001).

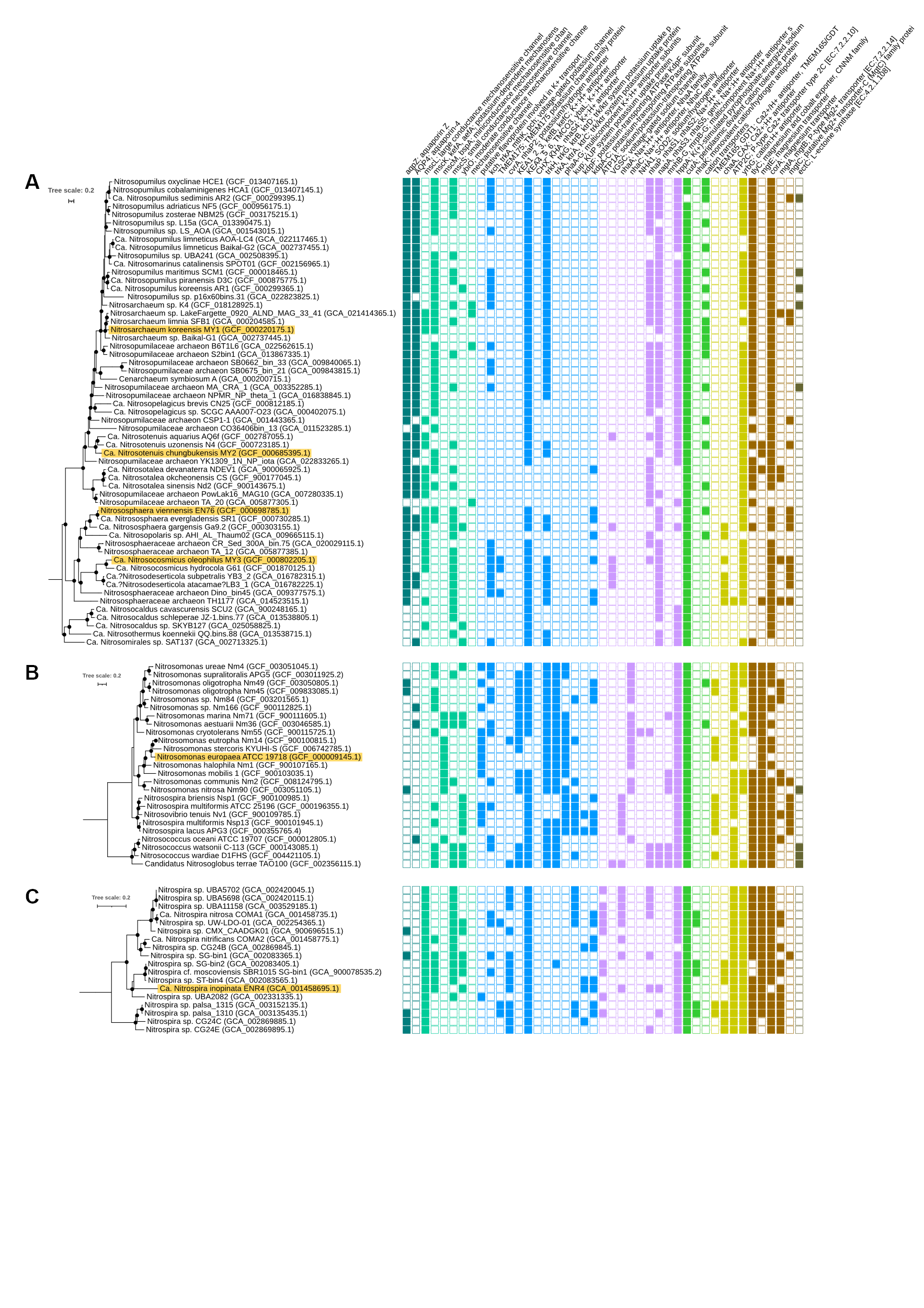

#### Fig. S5. Comparative genomic repertoire of osmoregulatory systems in ammonia oxidizers. The phylogenomic tree includes representative genomes of (A) ammonia-oxidizing archaea (AOA; *n*= 57), (B) ammonia-oxidizing bacteria (AOB; *n*= 25), (C) comammox clade A (CMX-A; *n*=14) and comammox clade B (CMX-B; *n*= 5). The complete dataset includes 463 genomes (see Dataset S1). The tree was constructed using the Anvi’o phylogenomics workflow (see Materials and Methods for details). Black circles indicate nodes with ≥70% bootstrap support. Strains used in salinity-dependent growth experiments (see Fig. 5) are highlighted in yellow. Gene presence or absence was determined based on KEGG annotations. See the Methods section for details on gene selection and annotation procedures. Solid and open squares represent gene presence and absence, respectively.

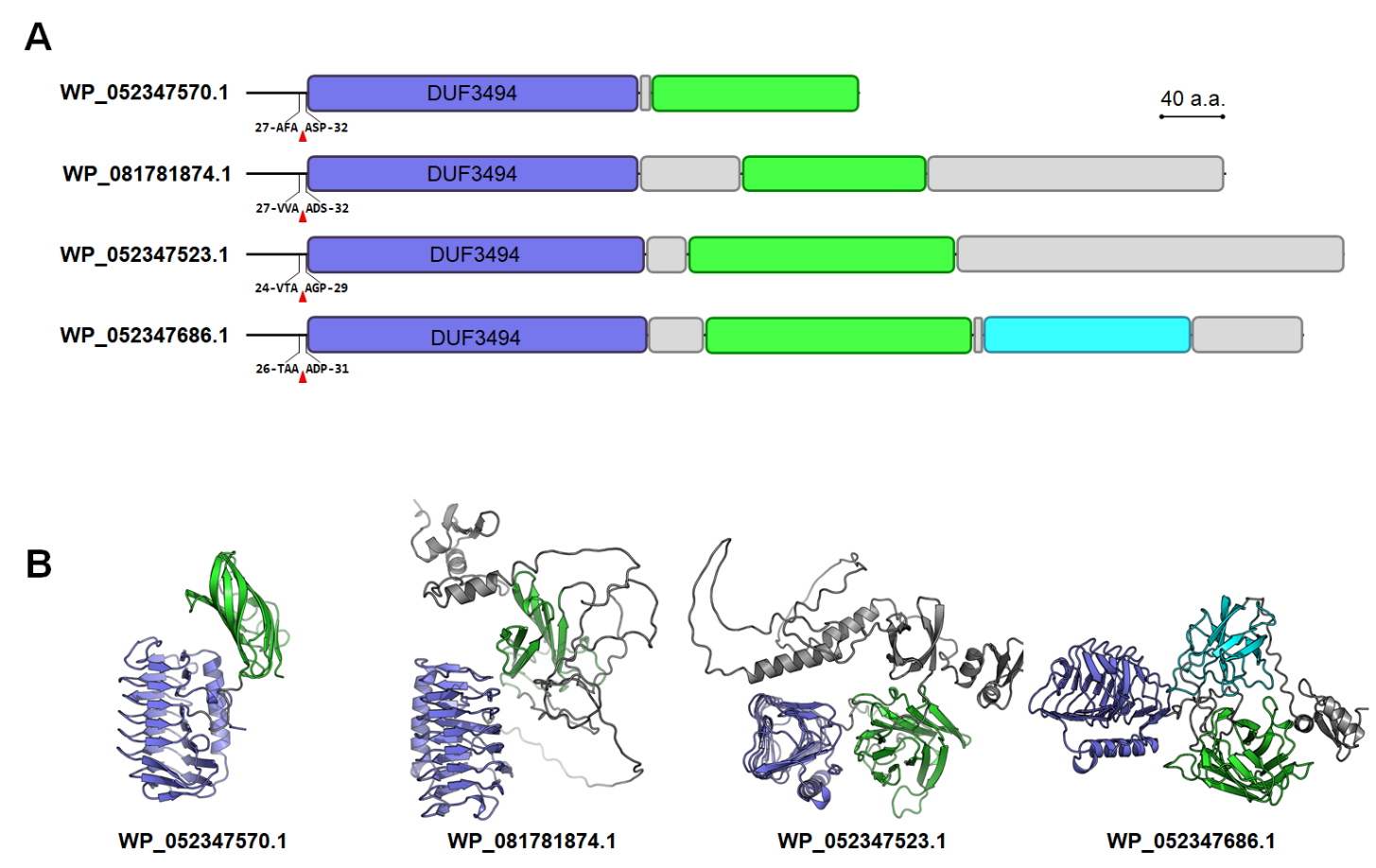

#### Fig. S6. Sequence analysis and structural prediction of IBP-like proteins from “*Candidatus* Nitrosotenuis chungbukensis” MY2. (A) Signal peptide cleavage sites are indicated with red triangles. Schematic domain structures include DUF3494 domain (purple), β-sandwich domain (blue), and β-barrel domain (cyan), and loop regions (grey). (B) Ribbon representations illustrate the overall structures of the mature proteins (excluding signal peptides). The color scheme used in A is applied to distinguish individual domains.

### Legends for Supplementary Datasets

Dataset S1. Genomic inventory of osmoregulatory genes in ammonia oxidizers.

Dataset S2. ASVs of ammonia oxidizers, including the *Nitrospira* lineage, amplified from soil slurry and lake microcosm samples.

Dataset S3. Gene list and transcriptomic abundances in “*Ca*. Nitrosotenuis chungbukensis” MY2 cells grown under 1× and 10× MWM.

Dataset S4. Gene list and transcriptomic abundances in *Nitrososphaera* *viennensis* EN76 cells grown under 1× and 10× MWM.

Dataset S5. Selected genomic functions of “*Ca*. Nitrosotenuis chungbukensis” MY2 and *Nitrososphaera* *viennensis* EN76 with corresponding transcriptomic responses and proteomic detection under hypoosmotic conditions.
